# Visualization and Analysis of Whole Depot Adipose Tissue Neural Innervation

**DOI:** 10.1101/788885

**Authors:** Jake W. Willows, Magdalena Blaszkiewicz, Amy Lamore, Samuel Borer, Amanda L. Dubois, Emma Garner, William P. Breeding, Karissa B. Tilbury, Andre Khalil, Kristy L. Townsend

**Affiliations:** School of Biology and Ecology, University of Maine, Orono ME; Graduate School of Biomedical Science and Engineering, University of Maine, Orono ME; School of Molecular and Biomedical Sciences, University of Maine, Orono ME; Department of Chemical and Biomedical Engineering, University of Maine, Orono ME; CompuMAINE Laboratory, University of Maine, Orono ME; Department of Neurological Surgery, The Ohio State University, Columbus OH

**Keywords:** White adipose tissue, innervation, neurovascular interaction, myelination, confocal microscopy, widefield microscopy, whole mount, maximum projection, clearing, mouse

## Abstract

Little is known about the diversity and function of adipose tissue nerves due, in part, to the inability to effectively visualize the various nerve subtypes residing within these tissues. The tools currently available for researchers to image and quantify adipose tissue innervation are limited and dependent on optical clearing techniques and light sheet microscopy. Here we present a method of tissue processing that uses mechanical force to compress tissue to decrease tissue thickness in the z-axis by expanding it in the x and y-axes whilst leaving cells intact. This has been combined with autofluorescence quenching techniques to permit intact whole tissues to be mounted on slides and imaged on any microscope, with a complementary means to perform whole tissue neurite density quantification. We have included examples of how this technique can be used to further our current knowledge of adipose-nerve communication by characterizing the nerves, nerve-subtypes, and neurovascular interactions within subcutaneous white adipose tissue in mice.

## Introduction

Historically overlooked as a location of diverse peripheral nerve innervation [1], the adipose organ was most prominently inspected for innervation in the mid 1960’s when sympathetic nerve fibers were visualized within brown adipose tissue (BAT) [2]. BAT nerves were later comprehensively investigated by the Bartness research group [3]. Energy expending BAT was the first adipose tissue to be identified as being highly innervated due to its important role in thermogenesis [4], which requires significant sympathetic nervous system (SNS) input and release of norepinephrine [2, 5, 6]. Sensory innervation has also been documented in BAT, particularly around vasculature, and has been proposed to play a role in lipolysis [7]. More recently white adipose tissue (WAT), associated more with energy storage, but which is a dynamic and plastic tissue capable of undergoing transformation during a cold-induced “browning” process, was demonstrated to be highly innervated by sympathetic [8, 9] and sensory nerves [5, 10], but not parasympathetic nerves [11].

In order to visualize adipose innervation, it had been common practice to slice adipose tissue into 7-10 μm thick sections and immunolabel for various neuronal markers [11–14], which include the sympathetic nerve activation marker tyrosine hydroxylase (TH); the rate-limiting enzyme for the synthesis of catecholamine neurotransmitters such as norepinephrine. Several important findings emerged from this practice, but it was not without limitations. Thin sections of tissue reduced peripheral nerves to puncta, leaving investigators unable to accurately determine arborization or the ability to quantify innervation across an intact tissue. Importantly, our laboratory has revealed a new map of adipose anatomy in the inguinal subcutaneous white adipose tissue (scWAT) depot, and we and others have demonstrated that the pattern of innervation in scWAT is heterogeneous across the tissue [15] similar to the heterogenous browning potential also observed [16]. This warrants a more comprehensive look at adipose innervation across an intact depot or more anatomically-specific regional changes that may occur with physiological or pathophysiological stimuli.

Adipose tissue is notoriously difficult to image. Tissue resident lipids scatter light, and lipofuscin (especially prominent in unhealthy adipose) autofluoresces. Adipose tissue is also home to a dense vascular network which includes arteries, veins, capillaries, and lymphatics; all of which are also highly autofluorescent. Adipose depots can also be quite large in comparison to other organs, especially with the increased adiposity of obesity. Metabolically healthy and lean mice can have inguinal scWAT depots that are several millimeters thick and this increases drastically in metabolically unhealthy mice. Because of this, combating tissue autofluorescence has been a necessity for researchers, especially in whole depot imaging.

Methods for optically clearing and delipidating whole organs for imaging have been developed, particularly for brains, which are full of light scattering lipids similar to adipose. These techniques are currently being applied to imaging adipose tissue and specifically imaging and quantifying innervation within whole adipose depots [15, 17–22]. While useful for getting crisp images of the large myelinated nerve bundles in adipose, this technique does have its drawbacks. The current methods typically rely on quantifying only a few small so-called ‘representative’ three-dimensional tissue sections, which can miss the heterogeneity and regional anatomy of the intact tissue. Until the regional variation in neurite density in scWAT [15, 20] is better understood, it cannot be accurately reduced to a few representative images or a small tissue block. Furthermore, some of the most effective methods of tissue clearing use caustic reagents which, for optimal results, require immersion of specialized objective lenses (ie: “BABB-safe”) in the clearing media for refractive index matching, and many of these shorten fluorescence lifespan and can be detrimental to endogenous fluorophores. This can be costly and prevents tissue from being mounted on slides for easy widefield epifluorescence imaging. Some clearing media can also solidify at room temperature which causes yet another hurdle for researchers. There is also the separate issue of obtaining homogenous staining through thick cleared tissue. Traditionally, poor antibody penetration has had to be combatted with drastically increased incubation times and more recently with stochastic electrotransport [23] and barometric pressure [24].

For these reasons we have developed an alternative approach to adipose tissue processing that uses mechanical pressure to homogenously compress entire adipose depots to reduce tissue thickness. Z-axis compression has been used for other tissues as a means of reducing tissue thickness to allow for faster image acquisition [25]. We aimed at developing a method that will be accessible for a wide population of laboratories to use, which is a whole mount imaging technique that does not require optical clearing or tissue sectioning and can be used with either standard widefield or confocal microscopy without the need for specialized BABB-safe objective lenses. This is accompanied by a simple option for quantification of adipose neurites using methods that we have made publicly available (see Methods). Widefield microscopy is sufficient for many qualitative analyses of entire adipose depots, but for precise colocalization and quantitative analysis confocal imaging is required which can pose a significant increase to image acquisition times (see Methods). However, the cost for light sheet microscopes can create a barrier to entry that we wish to ameliorate, allowing more research groups to investigate adipose innervation and the role of adipose nerves in physiology and pathophysiology.

Like all of the other methods mentioned above, ours also has some drawbacks and limitations, which is that the tissue depot must be compressed mechanically for it to be imaged effectively on both widefield and confocal microscopes. Otherwise, our technique presented here maintains an intact adipose depot and allows for detailed visualization of up to five fluorescent channels at a time, either from endogenous fluorescent mouse reporter lines or from fluorescent antibody immunostaining (both of which maintain fluorophore integrity, in this method).

We have tested this technique using two separate scWAT depots (inguinal and axillary) which have been suggested to demonstrate different responses to neurotrophic stimuli and browning potential [15]. This whole mount technique has been optimized for the pan-neuronal markers PGP9.5 [26, 27] and β3-tubulin [28, 29]; the sympathetic nerve marker, tyrosine hydroxylase (TH) [14]; markers for sensory innervation that included advillin (AVIL) [30, 31] and Nav 1.8 [32]; the myelination marker myelin protein zero (MPZ) [33]; and others. This technique has also been optimized for non-antibody based fluorescent labeling approaches such as nuclear labeling with DAPI and vascular labeling with Isolectin IB4 (IB4). This has allowed us to further our understanding of scWAT in mice by characterizing the innervation that exists within this tissue with greater scrutiny, demonstrating neurovascular interactions, parenchymal innervation, and neuroimmune interactions.

## Results and Discussion

### Development of the whole mount technique

We have tested numerous clearing techniques with scWAT and BAT (Suppl. Table 1) which included ScaleA2 [34], BABB [35, 36], CUBIC [37], CUBIC CB-perfusion [37], iDISCO [38], uDISCO [39], UbasM [40], and a sucrose gradient method [41]. Each clearing method had its own set of trade-offs, with some methods distorting tissue morphology, limiting fluorescence lifespan, and requiring costly objective lenses to image (summarized in Suppl. Table 1). Not all clearing techniques sufficiently clear adipose, as well. These pros and cons have been well documented in various other tissues [42–45]. To circumvent these issues, we have developed a method of whole tissue processing, outlined in figure 1a, that allowed for whole adipose depots to be imaged on any epifluorescence widefield microscope or confocal microscope without the need for immersion lenses or caustic clearing agents. Entire processing time for an intact adipose depot with a single directly conjugated antibody takes as little as 6-days. This is significantly shorter than the majority of adipose clearing methods (10-14days) and is comparable to the remainder.

**Figure 1:**
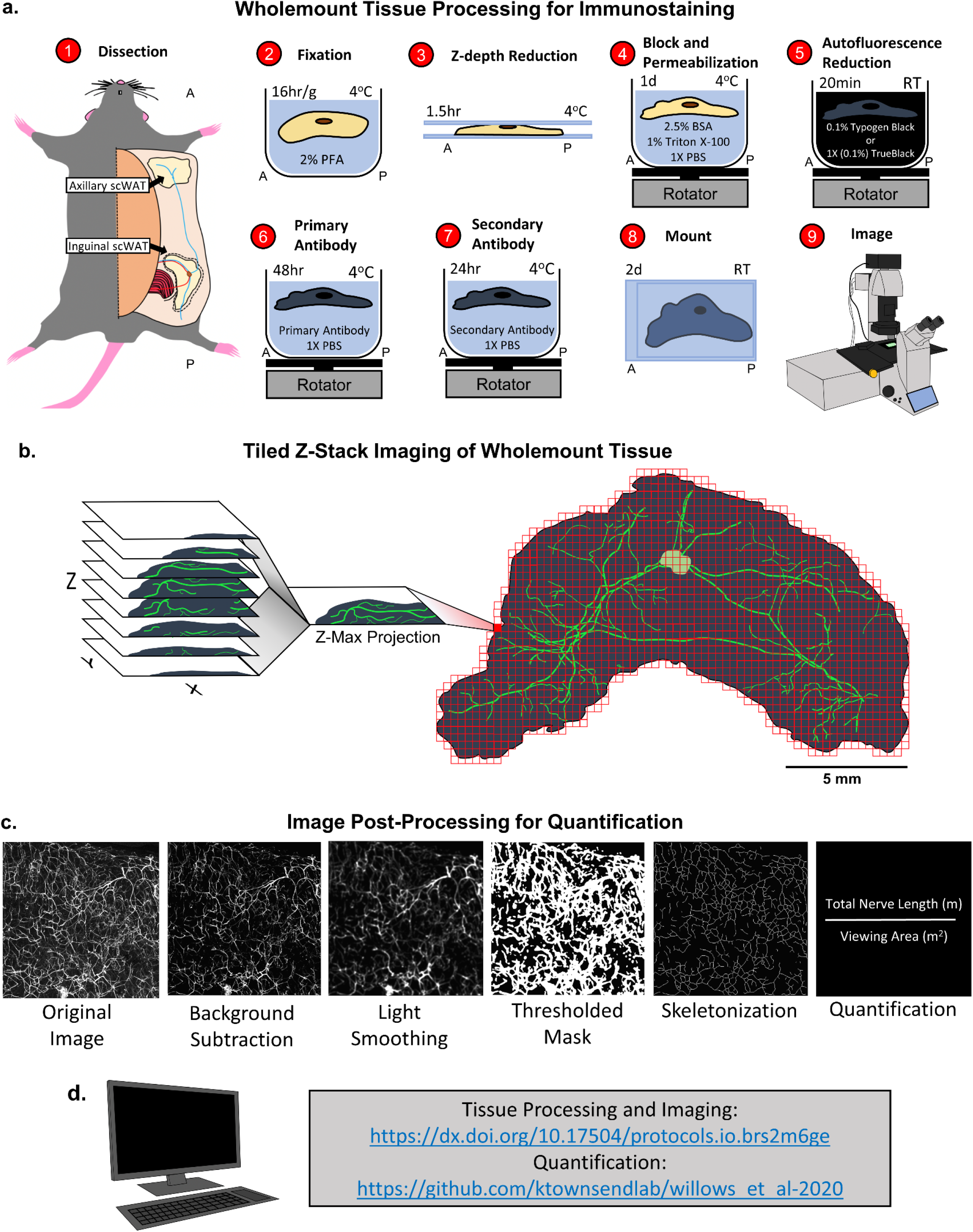
Adipose tissue processing, imaging, and quantification method. Diagram depicting steps of wholemount tissue processing technique (a). See methods for detailed description of each step. Tiled z-stacks were imaged for the entire tissue with either a 5X or 10X objective and a 2D maximum projection image was rendered (b). Post processing of single tile z-max projections for neurite density quantification (c). Each z-max projection tile was further processed by subtracting the background away and adding light smoothing. Next a thresholded mask was applied and the image was skeletonized. Total neurite length was calculated for each tile and averaged for the entire viewing area (c.) Link to whole mount protocols and script for quantification (d.)

This approach has allowed us to image entire tissue depots and construct 2D visualizations from 3D data acquisition by tiling z-max projection images taken at low magnifications using a laser scanning confocal microscope (Fig. 1b) and used for quantifying total nerve density (as described in Methods section) (Fig. 1c). Our quantification method is based on pixel intensity. By capturing data as a z-max projection we retain the highest pixel intensities from each z-plane to greatly decrease computation time for the quantitative analyses while keeping potentially lost data to a minimum. We have developed our quantification method using open access software and Matlab and all code has been made publicly available (Fig. 1d). To acquire a micrograph of an entire intact depot on a point scanning confocal microscope times are subject to the desired resolution, number of fluorescent markers, and size of the tissue. Acquisition time has averaged 13-16hrs when a single fluorophore is imaged at 10X objective magnification per tile at resolutions of 720×720 and a z-step sampling size of 12μm. We also experimented with confocal resonant scanning and subsequent denoising with Denoise.ai to reduce image acquisition time at the cost of resolution (Suppl. Fig 1). A whole depot was imaged in only 2hrs 35min (Suppl. Fig. S1a). Unfortunately, we found that by reducing resolution to image faster, we lost most of the small neurites when compared to the slower point scanning approach (Suppl. Fig S1b). Because of this we have determined that higher resolutions and longer imaging times are required to capture an accurate representation of parenchymal innervation.

Although the use of light sheet microscopy can decrease image acquisition time from hours to minutes, in our experience (using multiple platforms) imaging an intact whole adipose depot from various physiological states was extremely challenging with commercially available models. This led us to develop an adipose tissue processing technique that made imaging entire intact adipose depots (no need to bifurcate the depot) on a confocal microscope possible.

Whole mount processing of inguinal and axillary scWAT was achieved by first excising intact depots from the mouse and fixing in 2% PFA overnight. Tissue thickness was reduced to shorten incubation times, enhance antibody penetration, and reduce light scatter by flattening the fixed tissue between 2 large glass slides with binder clips to apply gentle pressure (Fig. 1a) in a process we have termed ‘Z-depth reduction’.

Tissues are then quenched for autofluorescence and mounted on glass slides with a glycerol based mountant. The glycerol based mountant acts, in part, as a clearing agent, though its use was dictated by the increased coverslip adherence the high viscosity provided. Mounting in a non-clearing aqueous based mountant offers the same quality images but we found that the coverslip adherence is poor, resulting in tissue oxidation and subsequent loss of signal, as well as increased autofluorescence.

To investigate how z-depth reduction altered tissue morphology we first measured the change in tissue area that occurred by z-depth reduction and subsequent mounting for axillary and inguinal scWAT depots (Fig. 2a). Z-depth reduction and mounting resulted in a significant increase to tissue area for both axillary (p=0.0006) and inguinal (p=0.0002) depots (Fig. 2b). In both cases XY area was increased approximately by a factor of three with tissue area increasing after z-depth reduction and again when mounted (Fig. 2b). Tissue thickness in the Z-dimension was measured at the thickest point of each tissue before and after Z-depth reduction. Tissue thickness was reduced significantly (axillary p=0.0230, inguinal p=0.0192) from approximately 2-3 mm before Z-depth reduction to less than 200 μm after mounting (Fig. 2c). It is important to specify that the tissues used here are from metabolically healthy young adult mice. Tissues from obese or lipodystrophic mice will likely follow the same trend, but exact changes may be different.

**Figure 2:**
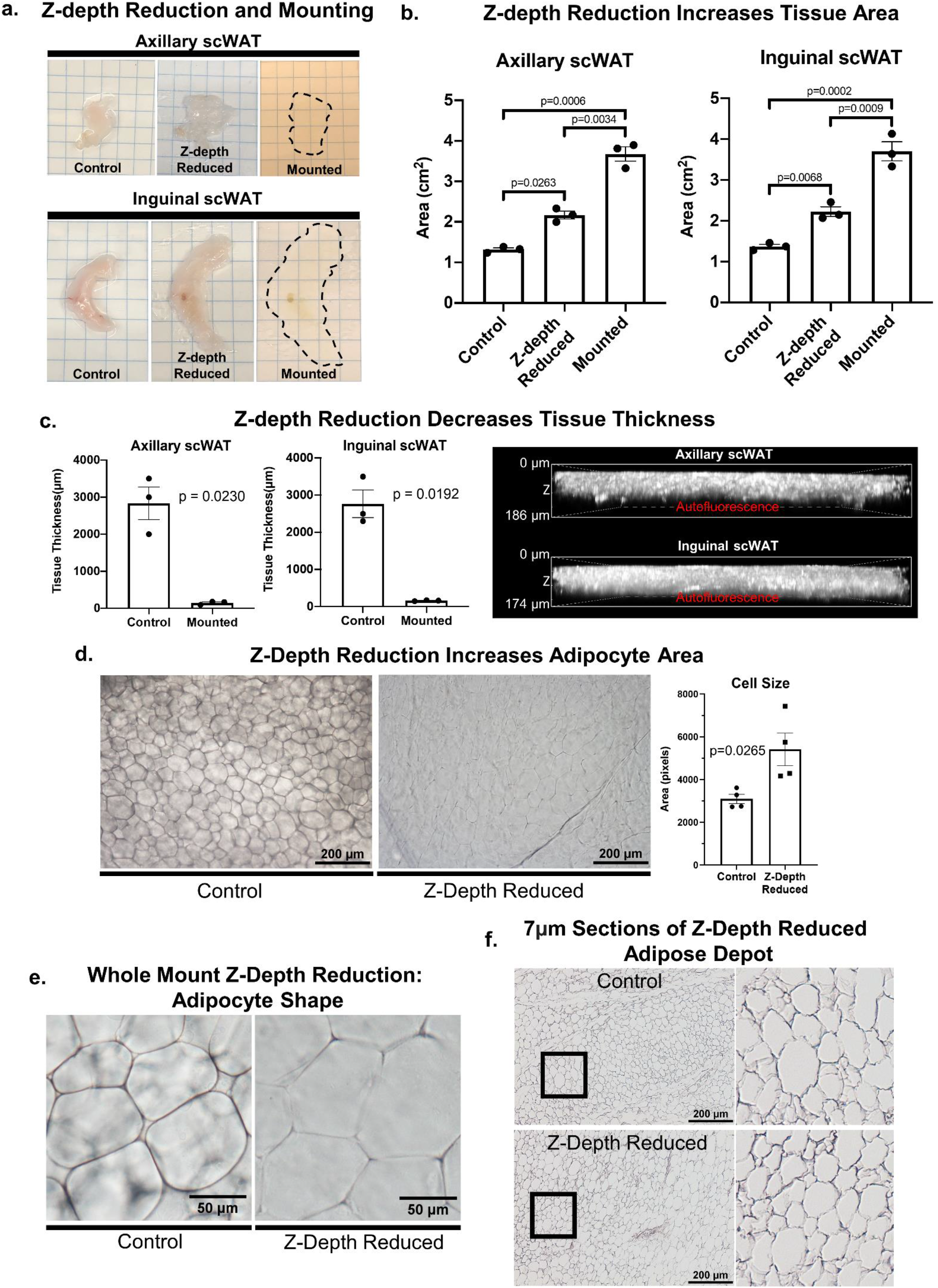
Z-depth reduction and tissue morphology. Axillary and inguinal scWAT depots excised from adipose were Z-depth reduced and mounted on slides (a). Tissue area was measured before Z-depth reduction, after z-depth reduction, and after mounting (b). N=3, paired two-tailed Student’s t-test, alpha level 0.05, error bars are SEMs (b). Tissue thickness in the Z-axis was measured at the thickest point before and after Z-depth reduction and mounting (c). Mounted tissues were measured by 3D projecting tissue autofluorescence captured on a Nikon A1R with a 20X objective (c). N=3, paired two-tailed Student’s t-test, alpha level 0.05, error bars are SEMs (c). Inguinal scWAT was either whole mount processed (Z-depth reduced) or left unaltered (control) and placed on a slide and imaged with transmitted light (d). Four representative 10X micrographs were captured for each tissue and the area for 30-120 cells was measured per micrograph and analyzed using a two-tailed Student’s t-test, Alpha level 0.05, error bars are SEMs (d). 3.5X Digital zoom of 10X transmitted light micrograph illustrating difference in cell shape (c). Hematoxylin staining of 7 μm inguinal scWAT sections that either received z-depth reduction before paraffin embedding or did not (control)(f). Transmitted light micrographs were captured on a Nikon E400 upright microscope (d-f). Scale bars are 186 μm and 174 μm (c), 200 μm (d,f) and 50 μm (e).

Next, we were interested in how the adipocyte size was affected by the increased tissue area, and whether cells remained intact. Because tissue area and thickness were altered similarly between both inguinal and axillary depots we narrowed our analysis to inguinal depots moving forward. Unstained whole inguinal scWAT depots were either Z-depth reduced and mounted or remained in their physiological state post-dissection and left unmounted. Transmitted light was used to visualize unstained cell structure (Fig. 2d). Tissue thickness was reduced by Z-depth reduction which increased light transmission and softened the cell outlines visible in the control tissues (Fig 2d). Cell size was averaged across four representative images of each tissue to quantify possible changes in cell size (30-120 cells counted per image). Only cells that had clear and in-focus boundaries (given this was not 3D imaging) were included in the quantification. The average cell area was increased in every instance (p=0.0265) (Fig 2d). Under closer inspection of individual cells, we found that intracellular gaps were reduced to make room for the expanding cell area and that cells, due to being pressed against one another, adopted a polygonal shape (Fig 2e). Hematoxylin staining of 7 μm thick inguinal scWAT sections showed that cells remained intact and structurally similar following z-depth reduction (Fig 2f). A caveat of this analysis is that we were only able to investigate cell integrity following Z-depth reduction and not tissue mounting, which was demonstrated to cause additional strain to scWAT depots.

Adipose depots are malleable by nature, especially the subcutaneous adipose that melds to changes in skin pressure, and thus this method of tissue processing takes advantage of that by increasing tissue area in the X and Y directions to reduce tissue thickness in the Z. This structural manipulation introduces slight changes to individual adipocytes but leaves cells intact, not dissimilar to natural changes caused by animal movements such as lying down or stretching. We believe the benefits of tissue penetration and imaging ease outweigh these drawbacks, however slight morphological changes to adipose using this approach cannot be ruled out.

Tissue autofluorescence has always been a significant problem when imaging adipose due to the inherent autofluorescecent nature of lipids and lipofuscin that reside within it [46]. Sudan Black B (henceforth referred to by its historic name Typogen Black) staining has been used as a treatment to quench tissue autofluorescence for decades in 7-10 μm thick sections [47]. We applied Typogen Black and two other autofluorescence quenching techniques, TrueBlack® and hydrogen peroxide (used in many optical clearing protocols [42]), and compared quenching abilities against fluorescence filter cubes spanning the visible spectra: DAPI (blue), GFP/FITC (green), Cy3/TRITC (orange), and Cy5 (far-red; purple) (Fig. 3a). All methods were able to reduce autofluorescence in the blue, green, and orange wavelengths similarly, but Typogen Black was the least effective. The far-red Cy5 channel posed interesting barriers. Though far-red excitation results in the least autofluorescence of unstained tissue, autofluorescence is intensified strongly by Typogen Black staining and was left unquenched by TrueBlack®. Hydrogen peroxide was the only effective means of quenching autofluorescence in all 4-channels (Fig 3a) but adds significant time to the overall protocol and could be utilized only if all channels are needed. We next wanted to see how 647nm fluorophores were affected by the various autofluorescence quenching techniques (Fig 3b). All methods appeared to reduce fluorophore brightness when compared to the control tissue. Hydrogen peroxide was the least masking, followed by Typogen Black, and then TrueBlack® which resulted in a very diminished signal respectively (Fig. 3b).

**Figure 3:**
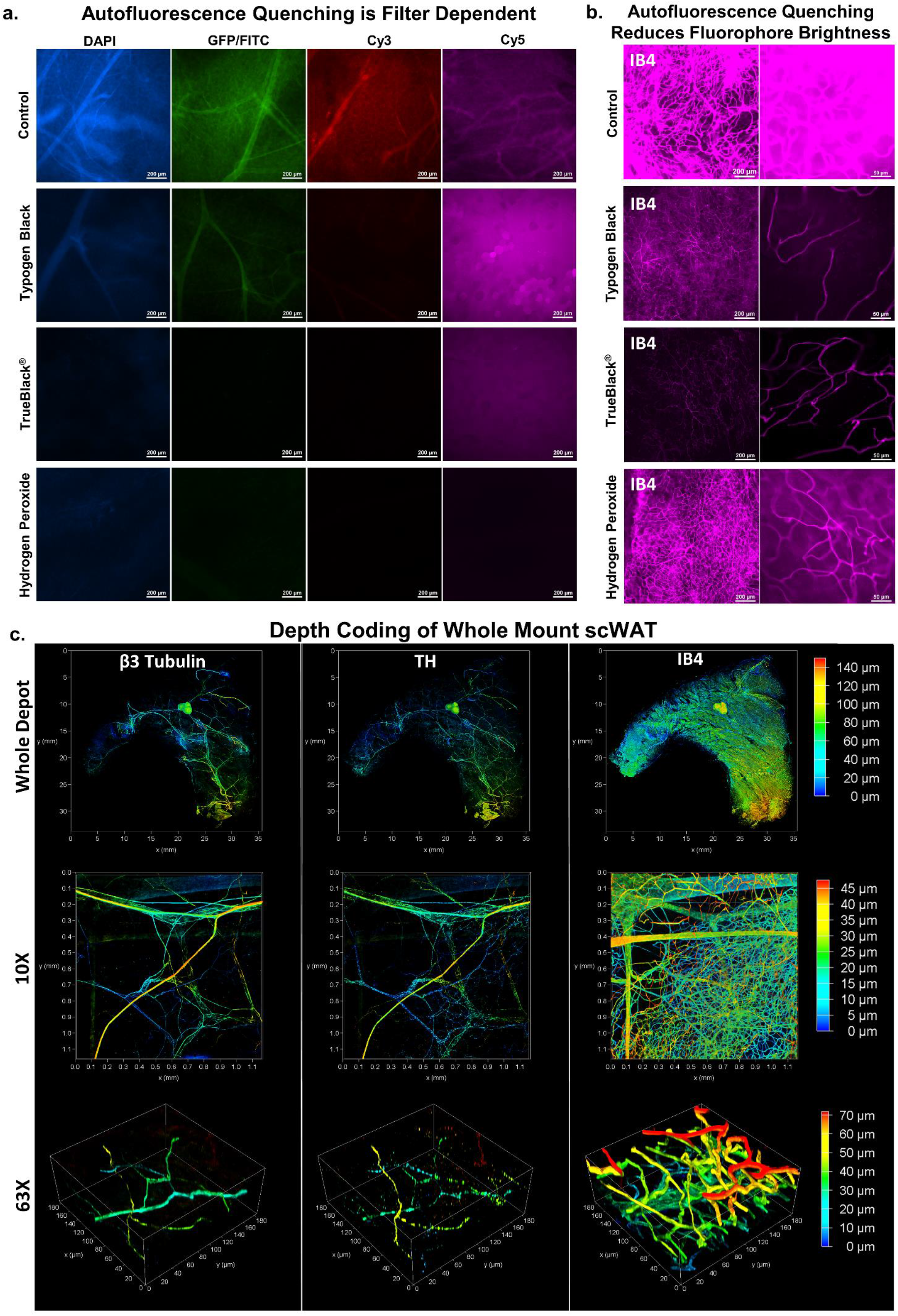
Whole mount adipose autofluorescence quenching and fluorescence staining. scWAT depots excised from *C57BL/6J* mice were whole mount processed and had autofluorescecne quenched with either Typogen Black, TrueBlack®, or 5% hydrogen peroxide. Autofluorescence was evaluated for each blocking method for 4 different fluorescent filter cubes spanning the visible spectra: DAPI, GFP/FITC, Cy3/TRITC, and Cy5 (a). Control tissue brightness was normalized in-post across all four filters and these settings were then applied to each image for a given filter: DAPI (used as control brightness), GFP/FITC (+10% brightness), Cy3 (+20% brightness), Cy5 (+40% brightness). Quenching methods were further compared for the Cy5 filter by staining the tissues with the vascular marker isolectin IB4 (IB4) conjugated to a 647nm fluorophore (b). Inguinal scWAT depot was whole mount processed and stained with β3-Tubulin, Tyrosine Hydroxylase (TH), and IB4 and depth coded (c). Whole depot was imaged as a tiled z-maximum projection with a 10X objective, (0-140um), a single tile captured at 10X (0-45um), and a high magnified section captured at 63X (0-70um) (c). Captured on Leica TCS SP8 DLS microscope (c). Scale bars are 200 μm (a), 200 μm and 50 μm (b). X and Y axes are 35 mm (c, whole mount), 1.1 mm (c, 10X) and 180 μm (c, 63X).

Although Typogen Black had the least autofluorescence quenching potential it was found to be sufficient for our uses and was preferred over TrueBlack®, which tended to mask true signal, and hydrogen peroxide, which increased processing times significantly. For these reasons Typogen Black was used when not staining with a 647nm fluorophore and TrueBlack® was used only when a 647nm fluorophore was used concurrently with other fluorophores.

Quenching dilutions were optimized in conjunction with pan-neuronal PGP9.5 staining to find the ideal balance between reduced tissue autofluorescence and undiminished true fluorescent signal (Suppl. Fig. S2a). Optimal concentrations, as determined by autofluorescence reduction and signal intensity, were found to be 0.1% Typogen Black (in 70% EtOH) and 1X or 0.1% TrueBlack® (2% TrueBlack® dye in 98% dimethyl formamide, diluted from 20X to 1X with 70% EtOH.)

An additional means of reducing tissue autofluorescence was performed by washing with 1X PBS with 10U/mL Heparin, which removed highly autofluorescent red blood cells (Suppl. Fig. S2b). Alternatively, perfusion can be performed to remove blood cells prior to tissue collection. Though the use of a glycerol-based mountant was decided upon due to its increased viscosity, it is also a known clearing agent [48], and we found that adding only a couple of drops to the tissue was enough to provide some level of optical clarity (Fig. 2a). This clearing was made more evident when tissues remained at room temperature for incubation and wash steps, prior to mounting, which resulted in reduced background autofluorescence (Suppl. Fig. S2c). It should be noted that autofluorescence when imaging with a widefield epifluorescent microscope is still quite present even after application of the aforementioned quenching techniques. Because of this, it is strongly recommended that confocal microscopy be used whenever possible to further reduce this background and non-specific signal.

To summarize: Typogen Black should be avoided when using far red excitation. TrueBlack® can be an acceptable replacement but it can mask fluorescent signal so avoid using with dim fluorophores. Hydrogen peroxide yields the best results all around but adds significant time to an already lengthy procedure. If only using one fluorophore it is best to use far-red excitation and do all washes and incubations at room temperature to avoid the use of autofluorescence quenching reagents entirely.

Secondary antibody non-specificity is likely to increase when immunostaining an entire tissue, and when primary-conjugated antibodies are not available or specific enough. We tested numerous secondary antibodies to confirm that we were only using those antibodies least likely to bind non-specifically (Suppl. Fig. S2d). Goat anti-rabbit IgG secondary antibodies, regardless of being highly cross adsorbed or not, did not result in significant levels of non-specific binding. Because goat anti-mouse secondary antibodies had such high levels of non-specific binding, we decided to avoid the use of all mouse primary antibodies for whole mount immunostaining moving forward.

Uniform staining of large whole mount tissues has always been problematic due to the increased incubation times required for antibodies to completely diffuse throughout the tissue [23, 49]. Incubation times were reduced by 1-2 days by reducing tissue thickness with z-depth reduction. Depth coding of immunostained scWAT following z-depth reduction indicated homogenous penetration of antibodies throughout the whole tissue (140μm total depth) (Fig. 3c). This was confirmed for various fluorescent staining approaches, including directly labeled β3-tubulin, indirectly labeled TH, and IB4 staining. Depth information can even be somewhat retained following z-maximum projecting by creating a height color coded projection, though no accompanying color-coded depth scale is provided at this time (Suppl. Fig. S3). Although we have not tried this ourselves, the addition of sucrose to 4% paraformaldehyde fixation can improve axonal staining by decreasing the appearance of “beading” along the axon [50].

### Characterization of peripheral nerves in scWAT

An inguinal scWAT depot was co-stained with two different pan-neuronal markers (PGP9.5 and β3-tubulin) and counterstained with DAPI to emphasize tissue cellular structure, and tiled images were z-max projected.

To demonstrate the versatility and capability of this technique we imaged tissues in two ways. Firstly, the whole adipose tissue was imaged at low magnification (5X) with limited resolution to observe overall morphology and fluorescence of the larger nerve bundles (Fig. 4a). Secondly, to visualize detailed innervation, areas of interest of the same tissue were imaged at 63X with much higher resolutions (Fig. 4b-c).

**Figure 4:**
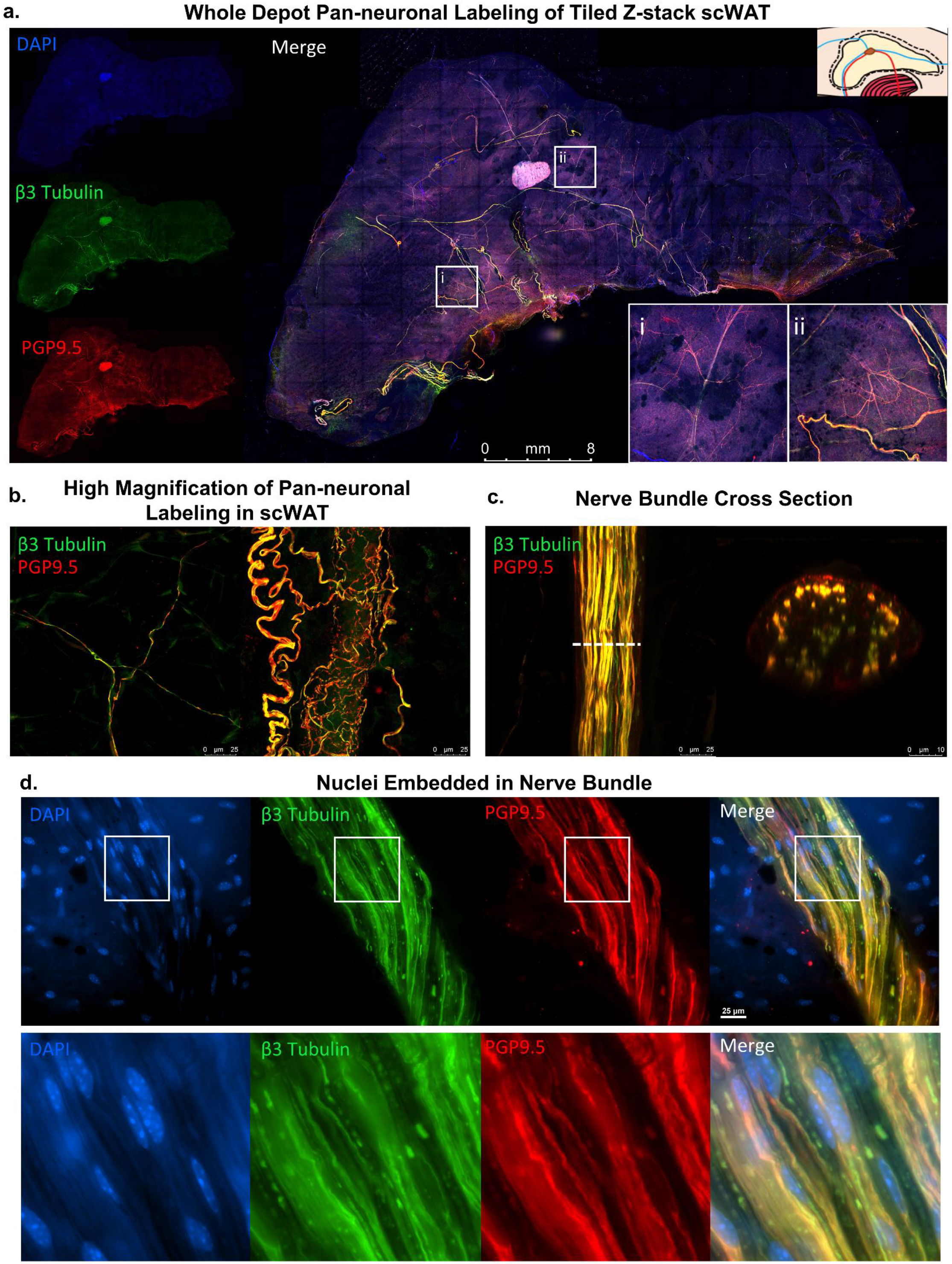
Peripheral innervation of scWAT. Inguinal subcutaneous depots from *C57BL/6J* mice were whole mount processed and stained with DAPI (blue) to show tissue morphology and the pan-neuronal markers β3-Tubulin (green) and PGP9.5 (red) captured on Leica TCS SP8 DLS microscope (a-c). Whole inguinal scWAT depot tiled z-max projections captured using 5X objective (a). White boxes show single tile expanded for visualization with 3.57X digital zoom-ins (i-ii). Representative images of small fiber innervation imaged as z-max projections with 63X objective (b). Nerve bundle imaged with 63X objective and digital cross sectioning performed with a 2.54X digital zoom (c). Nuclei on surface of nerve bundle and interlaced between nerve fibers captured in widefield on Nikon E400 with 60X objective (d). White box was digitally zoomed 3.9X (d). Scale bars are 8 mm (a) and 25 μm (b-d).

Whole tissue imaging exhibited near overlap of staining of the largest nerve bundles by both pan-neuronal markers (Fig. 4a). Nerves are unevenly distributed throughout the tissue in the X and Y-axis, and in the anterior-posterior anatomical directions. Because the tissues are compressed and subsequently z-max projected we were unable to accurately report nerve distribution within the z-axis. The highest concentration of large nerve bundles were present in the centermost third of the tissue (in the x-y direction), surrounding the subiliac lymph node (SiLN), as reported previously [15]. At greater magnification, z-max projection images showed an extensive network of neurites varying in diameter, running throughout the tissue and surrounding adipocytes as well as vasculature (Fig. 4b), also as previously noted[15]. Again, there appears to be almost complete fluorescent overlap of both pan-neuronal markers, with PGP9.5 being more readily visible on the smaller nerve fibers, which fits with the use of this antibody to assess small fiber neuropathy in the skin. Variation in immunofluorescence staining between the two pan-neuronal markers can be attributed in part to β3-tubulin being directly labeled and PGP9.5 being indirectly labeled (and thus, some off-target binding of the secondary antibody). Having a reporter mouse line instead of relying on antibodies would improve this technique. The innate autofluorescent quality of adipose tissue, though greatly reduced by Typogen Black or TrueBlack® staining, is still visible in most images and can help provide anatomical outlines. In fact, in most adipose clearing techniques in the published literature, autofluorescence of the tissue is exploited to reveal cellular architecture [20].

Digital cross section imaging of large nerve bundles demonstrated complete staining of the bundle but with gradation resulting in a weaker signal at the bundle’s center (Fig. 4c). Paraffin embedded cross sections do not show the same gradation (Suppl. Fig. S4), so we have attributed this observation to incomplete antibody penetration into the bundle. DAPI stained nuclei can be seen both surrounding the nerve bundle and residing between the numerous axons (Fig. 4d) (Suppl. Fig. S4). Since the nuclei of the neurons are located in the ganglia of the spine, the presence of nuclei within the nerve bundle suggests the presence of supporting cells (e.g. Schwann cells), immune cells, or potentially perineural adipocytes residing within the bundles.

Transmission electron microscopy (TEM) of scWAT sections revealed that putative nerves not only traverse through the tissue (Suppl. Fig. S5a) but may also come in direct contact with the adipocytes (Suppl. Fig. S5b-c). However, we have not yet confirmed synapses or tissue junctions directly onto adipocytes. Potential synapses have been observed in the stromal vascular fraction (SVF) of perigonadal adipose tissue (Suppl. Fig. S6a) as well as on blood vessels (Suppl. Fig. S6b), and on myeloid lineage SVF cells in inguinal scWAT (Suppl. Fig. 6c) using the post-synaptic marker PSD95. In general, the understanding of adipose tissue innervation has not extended to characterizing cellular interactions and whether they are synaptic (or a junction similar to the neuromuscular junction), or simply result from diffusion of neuropeptides and neurotransmitters from nearby free nerve endings.

To further characterize the nerves within scWAT, numerous immunostaining experiments were conducted by co-staining β3-tubulin with markers for either sympathetic nerves, sensory nerves, or myelination using the markers tyrosine hydroxylase (TH), advillin (AVIL), and myelin protein zero (MPZ) respectively. Whole mount imaging demonstrated fluorescence overlap of β3-tubulin and TH in the largest nerve bundles with extensive TH+ axons that spanned throughout the tissue (Fig. 5a). Nerve bundle digital cross sectioning showed that only some of the axons in each nerve bundle were TH+ (Fig. 5b). The small individual axons branching through the tissue, however, were nearly all TH+ (Fig. 5c), consistent with previous reports [11, 22].

**Figure 5:**
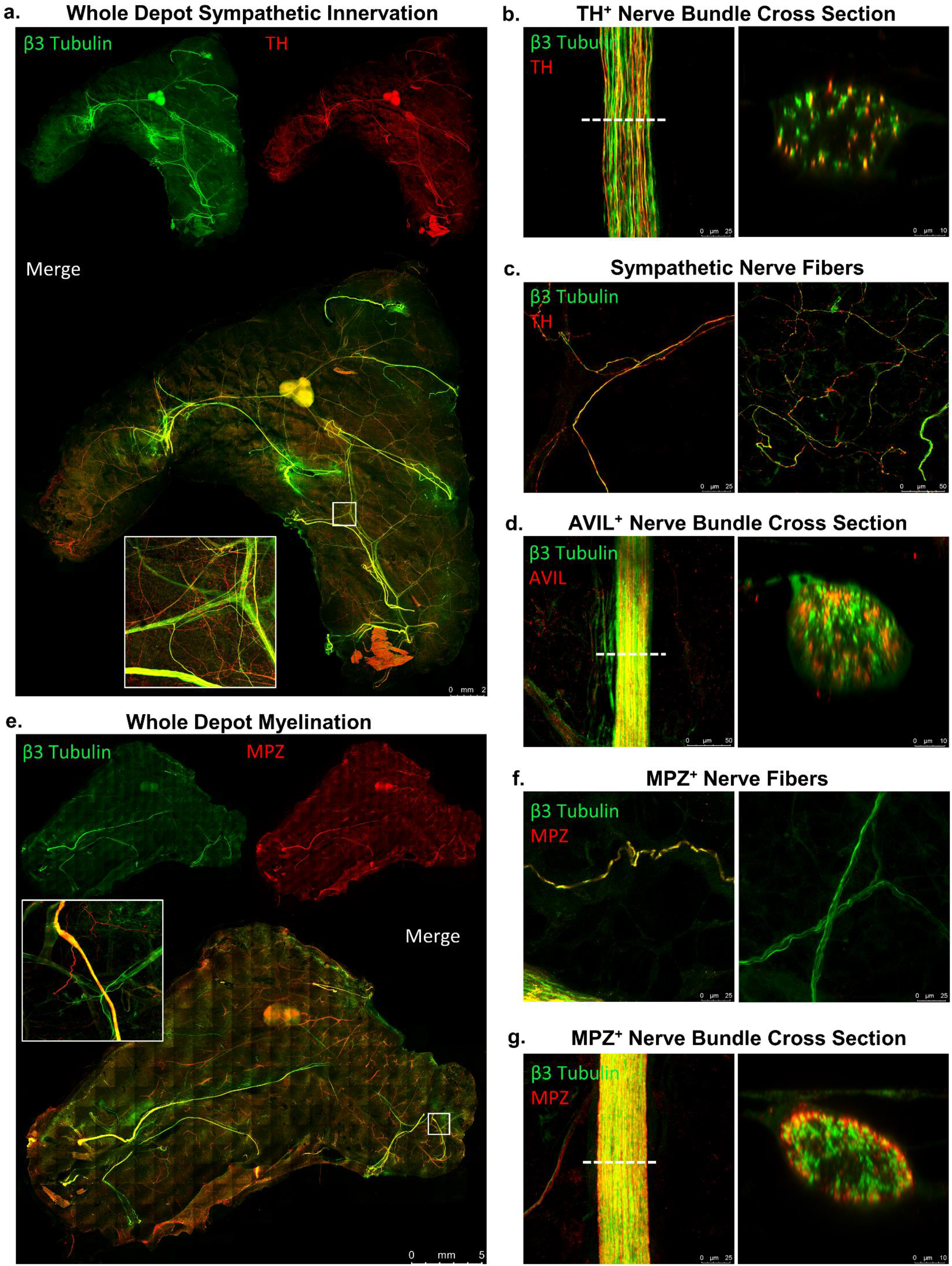
Characterization of whole mount scWAT nerves. scWAT depots from *C57BL/6J* mice were wholemount processed and stained with β3-Tubulin (green) and either TH (red) (a-c), AVIL (red) (d), or MPZ (red) (e-g). Whole mount tiled z-max projections of inguinal scWAT (a,d) and axillary scWAT (g) imaged with 10X objective. White boxes show a single tile expanded for visualization 5.75X (a,e). Nerve bundle cross sections imaged with 63X objective with a 2.54X digital zoom applied to the digital cross section rendering (b,e). Representative images of small fiber innervation imaged as z-max projections with 63X objective (c,f). All images captured on Leica TCS SP8 DLS confocal microscope (a-g). Scale bars are 2 mm (a), 25 μm and 10 μm (b), 25 μm and 50 μm (c), 50 μm and 10 μm (d), 5 mm (e), 25 μm (f), an 25 μm and 10 μm (g).

Digital cross sectioning of AVIL+ sensory nerve bundles revealed similar findings to cross sectioning TH+ bundles, with only some of the axons within bundles presenting as AVIL+ (Fig. 5d). This further suggested the presence of mixed nerves in adipose. It is worth noting that the AVIL antibody used here was observed to mark structural proteins in some large blood vessels as well, which accounted for the presence of AVIL+ regions that were β3-tubulin- when viewed as a tiled image. At greater magnification these large blood vessels did not interfere with imaging, and the lack of AVIL+ nerve fibers became apparent. Due to this, AVIL would appear to be a less-than-ideal marker for quantifying sensory innervation in conditions that may also alter vascularity. It is thought at this time that an AVIL reporter mouse or a Nav 1.8 reporter mouse would be a far superior method of labeling sensory axons.

Therefore, *Nav 1.8-Cre x tdTomato* reporter mice were also used to investigate the presence of sensory innervation in scWAT. Nav 1.8 marks sodium channels specific to sensory nerves [32] and fluorescence imaging showed a number of Nav 1.8+ sensory nerves throughout the tissue (Suppl. Fig. S7a). These were both large bundles and smaller parenchymal fibers, which complements our own results with AVIL imaging and current literature suggesting sensory innervation plays a significant role in WAT metabolic function [14, 51]. As observed with AVIL immunostaining, imaging of inguinal scWAT from *Nav 1.8-Cre x tdTomato* reporter mice also suggest the presence of mixed nerve bundles in this adipose depot (Suppl. Fig. S7a).

Whole mount staining with β3-tubulin and MPZ revealed that nerve bundles 25 μm and greater in diameter in scWAT are myelinated (Fig. 5e). In concordance with immunostaining, Luxol fast blue (myelin stain) staining of whole inguinal scWAT depots was performed which showed that the most highly myelinated nerves traverse through or nearby the SiLN (Suppl. Fig. S7b). It is not yet clear if these are mainly sensory axons that are myelinated, or a mix of sensory and sympathetic. Z-max projection imaging of parenchymal nerve fibers revealed that the majority of the smaller nerve endings are unmyelinated (Fig. 5f). Digital cross sectioning of MPZ+ nerve bundles showed that all of the axons within the bundle appeared myelinated, with the greatest fluorescence intensity being on the exterior of the bundle (Fig. 5g). At this time, it is thought that the gradation of the MPZ staining is caused by incomplete antibody penetrance but it cannot be ruled out that the exterior axons of these bundles are more myelinated than the interior axons. This potent MPZ+ staining on the bundle periphery also acts to mask some of the β3-tubulin+ nerves that reside within. TEM imaging was used to show both myelinated and unmyelinated putative neurites in contact with adipocytes (Suppl. Fig. S5b-c).

Our whole mount imaging technique has validated the presence of mixed bundle nerves residing in WAT as shown previously [17] and allowed for further investigation by creating digital cross sections of these mixed bundles. High magnification z-max projections revealed that the majority of neurites within the scWAT parenchyma are TH+, however, this was shown using TH-antibody which is known to be somewhat non-specific. Studies should be redone using a TH-reporter mouse to be certain. A relative minority of axons in the tissue are myelinated, and those that are reside mostly in large nerve bundles or run in close proximity, however, the contribution of myelinated versus unmyelinated axons to tissue function is currently unknown. Quantified colocalization of the various neuronal markers outlined above (PGP9.5, β3-tubulin, TH, MPZ, AVIL) at the whole tissue level and on nerve bundle cross sections can serve as another route for continued investigation.

### Neurovascular interaction in scWAT

The autonomic nervous system, comprised of sympathetic, parasympathetic, and sensory nerves, is required for regulating vascular tone [52] throughout the body, and WAT is no different. Generally, parasympathetic fibers are responsible for vasodilation and sympathetic nerve fibers are responsible for vasoconstriction, though this response is receptor specific and sympathetic nerves can mediate vasodilation as well [52]. It has been suggested that WAT lacks parasympathetic innervation [11] indicating that precise control of vasodilation is either not required in WAT or regulated entirely by sympathetic and sensory innervation. Neurovascular staining of whole mount scWAT was performed by co-staining with pan-neuronal markers and IB4, a marker for vasculature, as it binds to erythrocytes and endothelial cells [53–57] and effectively marks vessels smaller than 50 μm in diameter. Wholemount axillary scWAT tiled z-max projections exposed a dense vascular and lymphatic network residing in the scWAT depot (Fig. 6a). In general, the highest concentration of nerve bundles and blood vessels that are greater than 25 μm in diameter are located within close proximity of one another. These observations are consistent with what we have observed in inguinal scWAT [15]. Close inspection of the neurovascular interactions within scWAT revealed three specific and reoccurring types of interactions (Fig. 6b): i) Large nerve bundles and blood vessels that run next to each other within the tissue but do not seem to interact or intersect, ii) Large nerve bundles that have a vascular supply, likely providing nutrients; or a vasa nervorum, iii) Blood vessels that are highly innervated by smaller neurites which may regulate vasoconstriction; like a perivascular sympathetic plexus. In support of this, TEM imaging revealed a small putative axon in the stromal vascular fraction in between adipocytes in close proximity to a capillary (Suppl. Fig. S5a). Sympathetic innervation of scWAT blood vessels was also analyzed by co-staining β3-tubulin with TH and IB4. Small TH+ nerve fibers were found lining many of the vessels (Fig. 6c). Although small capillaries had significantly less innervation compared to larger arterioles, some TH+ nerves were found running along them. This supports the current literature that capillaries are relatively lacking in sympathetic innervation, whereas larger vessels have more sympathetic innervation [58–60]. Larger blood vessels likely also have sensory innervation as well [61].

Not all nerves within the tissue are lining blood vessels; many can be found branching the gaps from one blood vessel to another or disassociated from the blood vessels entirely. The vasa nervorum, or the blood vessels supplying nutrients to the nerve bundle, were digitally cross sectioned to show a blood vessel branching around a nerve bundle which contained 2 sympathetic axons (Fig. 6d). Extensive innervation of arterioles could be found surrounding both large and small diameter vessels (Fig. 76e). Of the nerves running throughout scWAT, the majority of MPZ+ nerves were found running along blood vessels and also had nuclei directly contacting them, as indicated by co-staining with DAPI, β3-tubulin, MPZ, S100β (a Schwann cell marker), and IB4 (Fig. 6f). Only one or two of the largest nerves lining each vessel tended to be myelinated. The smallest sympathetic projections that often engulfed many of the arterioles, when present, were found to be unmyelinated.

**Figure 6:**
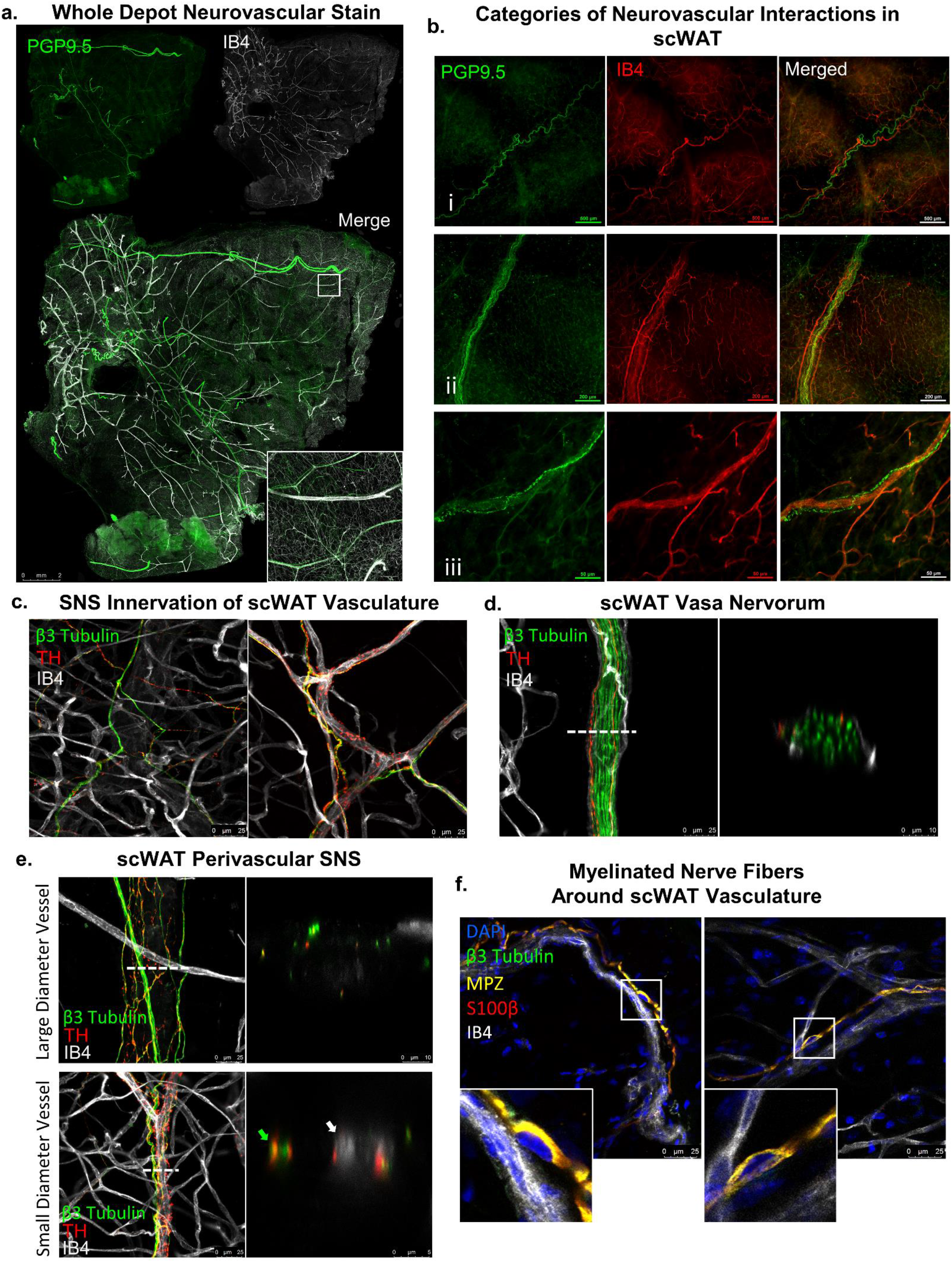
Neurovascular interaction within whole mount scWAT. Inguinal scWAT from *C57BL/6J* mice were wholemount processed. Whole axillary scWAT depot stained with PGP9.5 (green) and IB4 (white) was captured as tiled z-max projections imaged with 10X objective on Leica TCS SP8 DLS microscope (a). White box shows a single tile expanded for visualization 7.33X (a). PGP9.5 (green) and IB4 (red) used to display the 3 categories of neurovascular interaction within scWAT, imaged on Nikon E400 with 4X, 10X, and 40X objectives respectively (b). Representative images of the innervation of blood vessels imaged as z-max projections with 63X objective. β3-Tubulin (green), TH (red), IB4 (white) (b-d). Sympathetic innervation of small blood vessels (c). Vasa nervorum of inguinal scWAT nerve bundle imaged as z-max projection digitally cross sectioned with a 2.54X digital zoom (d). Large and small diameter blood vessel digital cross sectioning (e). 2.54X digital zoom applied to large diameter vessel cross section, 6.96X digital zoom applied to small diameter vessel cross section (e). Representative images of myelinated nerve fibers around vasculature imaged as z-max projections with 63X objective. Nuclei in direct contact of myelinated nerve bundles present in both images and white boxes expanded for visualization 3.67X (f). DAPI (blue), β3-Tubulin (green), MPZ (yellow), S100β (red), IB4 (white) (f). Scale bars are 2 mm (a), 500 μm (b-i), 200 μm (b-ii), 50 μm (b-iii), 25 μm (c), 25 μm and 10 μm (d), 25 μm, 10 μm, and 5 μm (e), and 25 μm (f).

It is important to note several caveats with IB4 staining of vasculature. Mouse blood vessel diameter ranges from 250μm to 4μm, depending on type of vessel. Veins encompass the upper limit, while microvasculature such as capillaries encompass the lower limit [62]. Large diameter blood vessels (>100um) often lose their erythrocytes during tissue preparation. However, erythrocytes tend to remain in the microvasculature. Because of this, the microvasculature is doubly stained (erythrocytes and endothelium) whereas arteries and veins present only with stained endothelium. This reduction in fluorescence, most prominent in the largest vasculature, can even be seen starting as small as 50 μm in diameter though these vessels are still readily visible (Fig. 6). Also, large sensory nerve bundles, immunostained for AVIL, an actin binding protein expressed specifically in somatosensory neurons [30, 31], tended to be co-stained with IB4. IB4 binding non-specifically to sensory neurons has been reported in the literature [63, 64]. However, only large sensory nerve bundles were observed to be labeled by IB4, eliminating the risk of misidentifying capillaries and, due to tissue morphology, positively stained nerve bundles can be easily distinguished from the vasculature.

### Whole mount imaging allows for quantification of total depot innervation and vascularity

Axillary scWAT depots taken from male C57BL/6J mice, housed at room temperature or cold exposed at 5°C for 7-days, were processed for whole mount imaging using the pan-neuronal marker PGP9.5 (Fig. 7a). The neurite densities per tile were then averaged for the entire area of the tissue for comparison among treatments. Quantification of images allowed for statistical analysis between experimental groups which showed that three of the four RT mice had less neurite density than all of the cold exposed mice but that overall this difference was not statistically significant (p=0.414) (Fig. 7b). This is similar to our previous studies using inguinal scWAT depots which used an alternative, but complementary, approach to whole tissue neurite density quantification [15]. Whole depot quantification might blunt regional differences in innervation, that may occur, for example, around the lymph node. Skeletonizing and quantifying neurite density from z-maximum projections, by definition, results in under sampling data. To verify the extent of this potential loss of spatial information, we calculated the neurite density on the individual z-slices that make up the 3D confocal data cube and then synthesized these quantities in two different ways: by taking the average neurite density score across the confocal stack (Suppl. Figure S8a), or by taking the maximum neurite density score across the confocal stack (Suppl. Figure S8b). In both cases, there is a very high correlation between the synthesized neurite density score (either obtained by averaging or by taking the maximum) and the neurite density score obtained from the z-max projection image (p<0.0001.) Of course, this does not allow us to conclude that no information is being lost by restricting the analysis to 2D z-max projections. However, it does allow us to conclude that the potential gain that one might make by considering all of the individual 2D slices instead of only the z-max projection would have to be explored beyond simple statistical measures such as taking the average or the maximum. And additionally, this potential gain in information would have to be weighed against the additional computational analysis time and effort. Neurite density per z-max tile was portrayed as a heatmap to better visualize region specific differences in neurite densities (Fig. 7c). This quantification and analysis approach can also be applied to vasculature (Fig. 7d).

**Figure 7:**
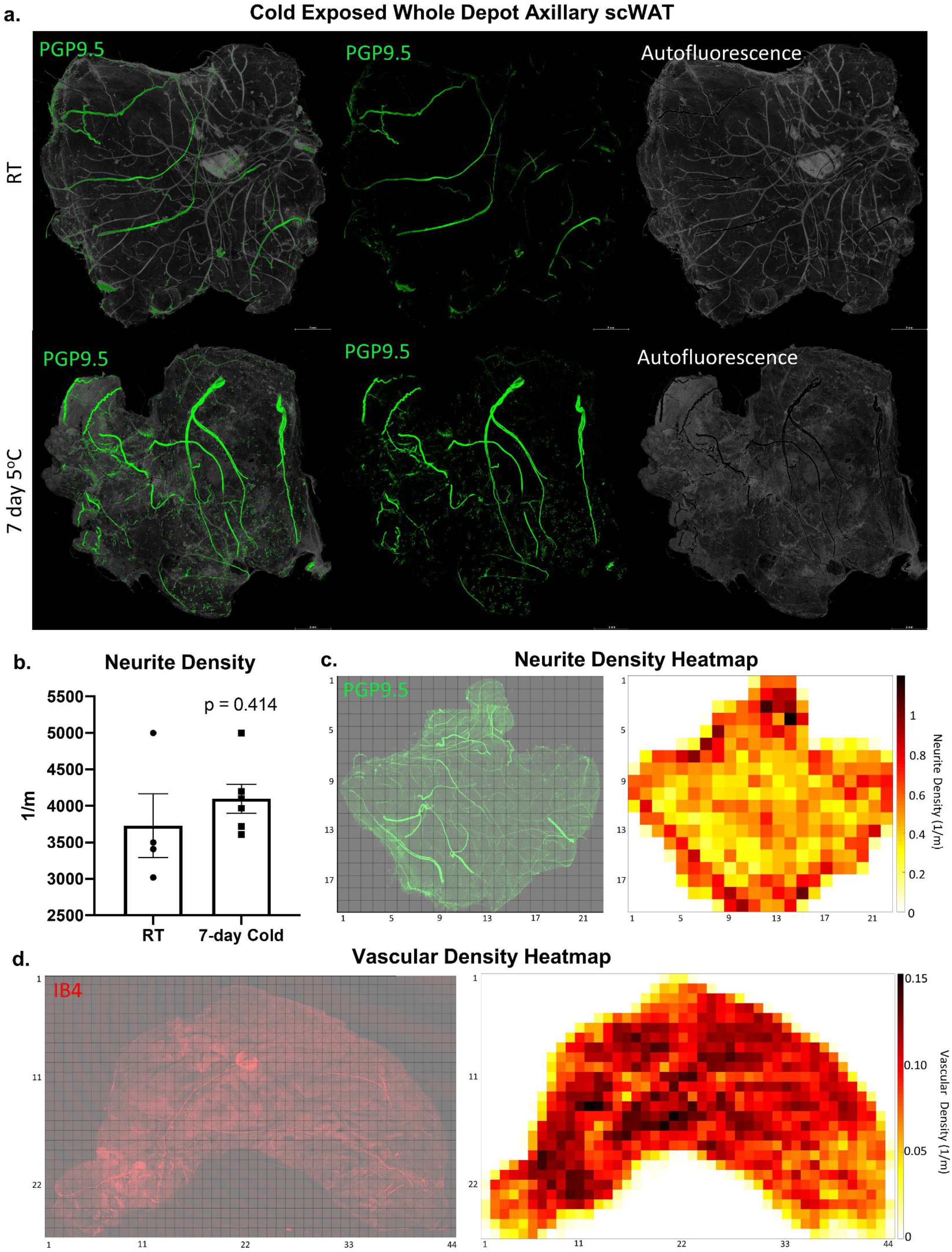
Whole depot neurite density quantification and heat map generation. Female *C57BL/6J* mice, aged 14-16wks, were either cold exposed (5°C) (n=6) or at room temperature (n=4) for 7 days. Axillary scWAT depots were wholemount processed and stained with PGP9.5 and imaged with 10X objective. To demonstrate the difference between true signal and tissue autofluorescence pixel intensity less than 55 was considered to be autofluorescence (grey) and pixel intensity 55 or greater was considered true staining (green) (a). Images were z-max projected, tiled, and thresholded and neurite density quantification was performed, compared across the cohort, and analyzed using a two-tailed Student’s t-test (b). Alpha level 0.05, error bars are SEMs (b). Neurite density per z-max tile was calculated and used to generate a heatmap (c). Inguinal scWAT depot was stained with IB4 to mark vasculature, imaged at 10X, and vascular density per z-max tile was calculated and used to generate heat map (d) Each tile is comprised of a 0.12cm × 0.12cm area (c-d). All imaging was performed with a Leica TCS SP8 DLS laser scanning confocal microscope. Scale bars are 2 mm (a).

### Second harmonic generation imaging

We also visualized the changes to scWAT innervation of morbidly obese mice which we have demonstrated to have adipose neuropathy (BTBR^*ob/ob*^); [15]. This proved technically difficult to accomplish using epifluorescent or standard confocal microscopy, largely due to the fibrotic nature of obese adipose tissue that led to severe autofluorescence that could not be mitigated with quenching. However, with second generation harmonic label-free imaging using 2-photon microscopy, it was possible to visualize nerves and collagen within scWAT of BTBR^*ob/ob*^ animals (Suppl. Fig. S9). While collagen is abundant in both BTBR^+/+^ and BTBR^*ob/ob*^ animals, a greater degree of colocalization and apparent alignment between collagen fibers and nerves is visible in BTBR^*ob/ob*^ animals (Suppl. Fig. S9), likely contributing to our inability to image the adipose nerves successfully in this mouse model.

## Conclusion

The whole mount processing and imaging technique presented here is meant to broaden the imaging capabilities currently available to researchers. We do not present a method meant to fully replace the need for optically clearing adipose depots or the use of light sheet microscopy but wish to provide an alternative method when such imaging techniques are not required or available. We have presented data to support that these more complex techniques are in fact not always required to get high quality images of adipose innervation. We have demonstrated how our method can be applied to imaging on affordable widefield epifluorescence microscopes as well as laser scanning confocal microscopes, to illustrate that this method of tissue processing can be applicable to researchers regardless of the tools they have available. The ease and accessibility of being able to take an entire adipose tissue and mount it on a slide to be imaged on any microscope cannot be overlooked.

Though the time to image an entire depot at resolutions sufficient for analyzing the smallest parenchymal neurites is substantial, this is only necessary when whole tissue quantifications are desired. For all other instances capturing a traditional single tile micrograph will take only seconds. Due to the broad adaptability of this technique, it is even possible to analyze gross tissue morphology and aberrant changes in the entire tissue in only a couple of hours, if desired, by lowering magnifications, decreasing resolutions, or by using a resonant scanning microscope.

The data collected herein reveal a rich and diverse neural innervation in mouse inguinal scWAT, likely indicating physiological roles that are yet to be uncovered. For example, differences in myelinated vs unmyelinated axons, heterogeneity of TH+ nerves, and the contribution of sensory nerves in adipose are yet to be clarified. In addition, the localization of true synapses or tissue junctions, versus release of nerve products from free nerve endings, and whether these exist on adipocytes or cells in the stromal vascular fraction (SVF) of the tissue, will also be important to uncover. The availability of new techniques allowing nerve visualization under various physiological and pathophysiological conditions will help the field advance an understanding of adipose innervation and brain-adipose communication, including neurovascular and neuroimmune contributions, and plasticity of neurites in adipose.

We have recently demonstrated diminished innervation of adipose tissue under circumstances of obesity, diabetes, and aging, a condition we have called ‘adipose neuropathy’ [15]. Neuropathy of adipose tissue (ie: a reduction in neurite density that may represent pathological nerve die-back) could have effects beyond the removal of sympathetic nerves releasing norepinephrine. Given that other nerve products, including from adipose tissue sensory nerves (ex: Substance P, VIP, CGRP, etc.) are found in adipose depots, the loss of proper innervation could also impact the physiological contributions of these neuromodulatory substances in adipose. Whether or not adipose sensory nerves directly communicate fuel status to the brain, perhaps in complement to endocrine factors such as leptin, is still an open question. Incoming sympathetic nerves may release norepinephrine in order to affect vasoconstriction, and also to impact local adipocytes and immune cells via synaptic and non-synaptic connections. We hope that by providing our method as a resource for investigators that these questions can be answered.

## Acknowledgements

Dr. Benjamin Harrison of UNE for developing earlier iterations of whole depot innervation quantification method (published in Blaszkiewicz et al. 2018 PLOS One). Dr. Clarissa Henry of UMaine for Leica light sheet confocal microscope training. Morganne Robinson, Thomas Szewczyk and James Miller for technical assistance. Dr. Ian Meng of The University of New England for gifting *Na_v_1.8* reporter mice adipose tissues. Dr. Dustin Updike of MDI Biological Laboratory for assistance with pilot imaging studies. Dr. Seth Tyler and Kelly Edwards of UMaine for TEM technical assistance.

## Author Contributions

JWW wrote the manuscript, created and optimized protocols, designed experiments, and conducted the majority of the microscopy studies and technical troubleshooting/optimization. MB wrote the manuscript, designed experiments, and optimized protocols. EG conducted TEM experiments. AL and ALD conducted clearing experiments. SB and AK developed neurite density quantification methods. WPB and KBT conducted 2-photon imaging. KLT wrote the manuscript, designed experiments, analyzed data, and oversaw the project.

## STAR Methods

### Animals

The following mouse strains were obtained from The Jackson Laboratory: C57BL/6J (Stock # 000664); BTBR^*+/+*^ (WT) and BTBR^*ob/ob*^ (MUT) (BTBR.Cg-*Lep^ob^*/WiscJ, Stock # 004824); LysM^cre/-^ (B6.129P2-Lyz2/J, Stock # 004781); BDNF^fl/fl^ (Bdnf^tm3Jae^/J, Stock # 004339); R26R-EYFP (B6.129X1-Gt(ROSA)26Sortm1(EYFP)Cos/J, Stock # 006148); TH^cre/-^ (B6.Cg-7630403G23RikTg(Th-cre)1Tmd/J, Stock # 008601. Myeloid lineage reporter mice were generated by crossing LysM^cre^x BDNF^flfl^ x R26R-EYFP. Nav1.8 reporter mice *(Nav 1.8^cre^ x tdTomato*) were a kind gift from Dr. Ian Meng, (University of New England, Biddeford, ME, USA). Animals were housed 3-5 to a cage (with the exception of BTBR WT and MUT animals which were housed in pairs), in a monitored temperature and humidity-controlled environment with 12/12hr light/dark cycle. Cages were cleaned weekly, ad libitum access to standard chow diet and water was maintained. For all studies animals were sacrificed using CO2 followed by cervical dislocation.

Adult male and female C57BL/6J were used for majority of immunostaining studies. Adult male BTBR WT and MUT were used in second harmonic generation imaging. Adult female TH-Rosa-eYFP reporter mice were used in optical clearing, and transmission electron microscopy studies. Adult male and female myeloid lineage reporter mice were used in high magnification imaging of adipose SVF. Adult female Nav1.8 reporter mice were used for whole mount immunostaining with PGP9.5 (GFP).

### Cold exposure experiments

All cold exposure was carried out in a diurnal incubator (Caron, Marietta, OH, USA) at 5°C. Adult male C67BL/6J mice were housed two to a cage and either maintained at room temperature or continuously cold exposed for 7 days. Inguinal and/or axillary scWAT was collected for wholemount tissue processing.

### Mouse adipose tissue collection and processing for immunofluorescence

Mice were euthanized using CO2 followed by cervical dislocation. Whole scWAT depots were carefully removed to remain fully intact and immediately fixed in 2% PFA at 4°C for 4hr-16hrs depending on thickness of tissue. The tissues were then rinsed for 30 minutes with 1X PBS w/ 10U/mL heparin, twice at 4°C. Tissues next underwent z-depth reduction in which tissues were flattened by being placed between two large glass slides bound tightly together with large binder clips, for 1.5hrs at 4°C. Tissues were incubated in blocking buffer (1XPBS/2.5% BSA/1% Triton X-100) at 4°C overnight but can be incubated up to 7-days if desired with blocking buffer replaced every 24hrs. After blocking period tissues underwent one of three methods of autofluorescence quenching: 1) Incubation in 0.1% Typogen Black for 20 minutes at room temperature on a rotator, 2) incubation in 1X (0.1%) TrueBlack® Lipofuscin Autofluorescence Quencher for 10 minutes at room temperature on a rotator, or 3) stepwise dehydration and subsequent bleaching with 5% hydrogen peroxide in 20% DMSO/MeOH as described by *Renier et al. 2014* [38]. Following quenching the tissues were washed with 1X PBS w/ 10U/mL heparin on rotating platform at 4°C replacing PBS every 1hr for a total of 4-6hrs, or until all unbound stain was removed. Tissues were incubated with primary antibody for 48hrs at 4°C. The following day tissues were washed with 1XPBS on a rotating platform at 4°C, replacing PBS every 1hr for a total of 4-6hrs followed by incubation with secondary fluorescent antibodies overnight. Tissues were then again washed with 1XPBS on a rotating platform at 4°C, replacing PBS every 1hr for a total of 4-6hrs. Following the immunostaining steps, when applicable, tissues were incubated with 1 μg/mL isolectin IB4 (IB4) (ThermoFischer, cat#:I32450 or cat#:I21413) diluted in HEPES buffer (pH 7.4), overnight at room temperature on a rotator. Tissues were then washed with 1XPBS on a rotating platform at room temperature for one hour twice. At this time, if DAPI co-staining was desired, the tissues were incubated in 100ng/mL DAPI for 10 minutes at room temperature on a rotator. Tissues then received four 1hr washes in 1XPBS at room temperature. At the conclusion of washing steps tissues were placed on large glass slides medial side facing up. A few drops of glycerol based mounting media was added to the tissue and coverslip was placed on top. Glycerol based mounting media was used because aqueous based mounting fluid lacks the required viscosity to fully adhere the coverslip when dealing with whole mount adipose. Slides were weighted down for 2 days and then sealed. Protocols for tissue dissection, processing for immunofluorescence, and confocal imaging can be found at https://dx.doi.org/10.17504/protocols.io.brs2m6ge.

### Widefield microscopy

All widefield micrographs were captured on a Nikon E400 with either a DS-fi2 color camera (Nikon, Japan) or Hamamatsu ORCA-Flash4.0 V2 Digital CMOS monochrome camera (Hamamatsu Photonics K.K., C11440-22CU) in conjunction with filter cubes for DAPI, GFP/FITC, Cy3/TRITC, and Cy5. Images captured on with Hamamatsu ORCA-Flash4.0 V2 were pseudo colored with NIS-Elements software (Nikon, Japan.) The following objectives were used: Nikon CFI Plan Apochromat Lambda 4X (NA 0.20, WD 20.0mm, dry), Nikon CFI Plan Apochromat Lambda 10X (NA 0.45, WD 4.00mm, dry), Nikon CFI Plan Fluor 20X (NA 0.50, WD 2.10mm, dry) and Nikon CFI Plan Fluor 40X (NA 0.75, WD 0.66mm, dry.)

### Confocal microscopy

All confocal micrographs were captured on a Leica TCS SP8 DLS (Leica Microsystems, Wetzler, Germany) laser scanning confocal microscope. Fluorescent labels were excited with either a diode 405nm laser (DAPI) or a white light laser (Alexa488, GFP, eYFP, Alexa532, Alexa555, tdTomato, Alexa 568, Alexa647) with excitation and emission spectra tuned specifically for each fluorophore or groups of fluorophores to reduce and eliminate spectral overlap when possible. All imaging was performed sequentially by frame. Photons were captured with both PMT and HyD detectors. The following objectives were used: HC PL FLUOTAR 5X (NA 0.15, WD 13.70mm, dry), HC PL APO CS2 10X (NA 0.40, WD 2.74mm, dry), HC PL APO CS2 20X (NA 0.75, WD 0.62mm, dry) and HC PL APO CS2 63X (NA 1.40, WD 0.14mm, oil.) Intact scWAT depots were imaged by tiling z-stacks across the full depth and area of the tissue. Whole depot micrographs in this manuscript were scanned bidirectionally at either 400Hz or 600Hz at 720×720 resolutions. Line averages ranged between 3-8 and z-step size ranged between 5-16 μm. Tissue thickness, though variable, was easily captured within the z-step range of the software (±250 μm of the focal point.) Identical image acquisition settings were applied for all tissues within cohorts that received neurite density quantifications. Anywhere between 4,000-65,000 individual images were captured per tissue which ranged from 100-900 tiles. These tiles were then merged together and processed into a 2D maximum intensity projection imaged either in LASX (Leica imaging software) or in Fiji [65] if being used for quantification. Whole depot image insets consisted of randomly chosen 10X z-max tiles of the merged image expanded for improved visualization. Representative images were captured at 1024×1024 resolution using a 63X objective and were z-max projected. Digital nerve bundle cross sectioning was performed by using the XZY scanning mode in the LASX software which applied a 2.54X digital zoom automatically to each image. All images were pseudo colored in LASX software.

### Neurite density quantification and heat map generation

Maximum intensity projections through the z-dimension (z-max projection) were generated for each tile individually using Fiji [65]. All single tile z-max projections were further processed using MATLAB x64 software (version 2018b, MathWorks). To remove low-frequency background noise, a 2D Gaussian smoothing kernel was used to convolve each z-max projection with a very large Gaussian blur using the *imgaussfilt* MATLAB command, with large standard deviation (~150), which was then subtracted from its corresponding original z-max projection. After background subtraction, a small gaussian blur (*imgaussfilt* with small standard deviation, ~2-3) was used to broaden out the neurite signal slightly before thresholding the image. Next, a binary thresholded mask was generated from the processed image. In the mask, only regions having an area larger than 40 pixels were kept for further analysis (using the *bwareaopen* MATLAB command). Then the *bwskel* MATLAB command was used to perform the skeletonizing procedure, with the added feature of removing any branches less than 4 pixels long. Except for an experimental analysis of 2D z-max projections vs. the individual 2D slices making up the 3D confocal data (Suppl. Fig S8), total nerve length was calculated using values measured for each 2D z-max projection. Total nerve arborization density was calculated as the ratio of total nerve length divided by the total viewing area, resulting in nerve length per square meter of tissue.

To generate heat maps a csv file was generated with the topological positions and the associated arborization quantity of each tile using the HeatMapChart function. Heatmap generation has multiple variables controlled by the user that could be explored further, such as the size of the subregion size and whether or not (and to what extent) the subregions may overlap. Code and instruction can be found at https://github.com/ktownsendlab/willows_et_al-2020.

It is also worth noting that the Python and MATLAB scripts developed for this study, provided above, allow the user to adjust the described parameters to account for batch differences due to variation in fluorescence intensity and autofluorescence. These adjustments can be made *a priori* or while looking at the skeletonized product of a single tile to quickly visualize how each adjustment changes the skeletonization.

### Antibodies and Fluorescing Stains

#### Primary antibodies included

PGP9.5 (1:1000, Abcam, Cambridge, U.K. Cat. #108986 and #10404); PSD95 (1:1000, Abcam, Cambridge, U.K. Cat. #ab18258); MPZ (1:250, Abcam, Cambridge, U.K. Cat. # ab31851); Advillin (1:500, Abcam, Cambridge, U.K. Cat. # ab72210); Perilipin-1 (1:100, Abcam, Cambridge, U.K. Cat. #ab61682); TH (1:250, Millipore, Burlington, MA USA; Cat. # AB152); neurofilament-M (2H3, 1:500) and synaptic vesicles (SV2, 1:250) from Developmental Studies Hybridoma Bank, (University of Iowa, USA); β3-Tubulin conj. Alexa Fluor 488 (1:250, Abcam, Cambridge, U.K. Cat. # ab195879); S100β conj. Alexa Fluor 647 (1:250, Abcam, Cambridge, U.K. Cat. # ab196175).

#### Secondary antibodies included

Goat anti-Rabbit IgG (H+L) Alexa Fluor 488 (1:500, ThermoFisher Scientific, Waltham, MA, USA; Cat. # A-11008); Goat anti-Mouse IgG (H+L) Alexa Fluor 488 (1:500, ThermoFisher Scientific, Waltham, MA, USA; Cat. # A-11001); Donkey anti-Goat IgG (H+L) Alexa Fluor 488 (1:500, ThermoFischer Scientific, Waltham, MA, USA; Cat. #A-11055). Goat anti-Rabbit IgG (H+L) Alexa Fluor 532 (1:500, ThermoFisher Scientific, Waltham, MA, USA; Cat. #A-11009); Goat anti-Rabbit IgG (H+L) Alexa Fluor 555 (1:500 or 1:1000, ThermoFisher Scientific, Waltham, MA, USA; Cat. #A-21428); Goat anti-Rabbit IgG (H+L) Alexa Fluor 594 (1:1000, ThermoFisher Scientific, Waltham, MA, USA; Cat. # A-11005); Goat anti-Rabbit IgG (H+L) Alexa Fluor 647 (1:1000, ThermoFisher Scientific, Waltham, MA, USA; Cat. # A-11012).

#### Fluorescent stains include

For vascular staining, tissues were incubated in isolectin IB4 stain conjugated to either Alexa Fluor 594 (1ug/ml, ThermoFisher Scientific, Waltham, MA, USA; Cat. # I21413) or 647 (1ug/ml, ThermoFisher Scientific, Waltham, MA, USA; Cat. #I32450). For nuclear staining, tissues were incubated in DAPI (100ng/ml, Sigma-Aldrich, St. Louis, MO, USA; Cat. # D9564).

### Luxol Fast Blue Myelin Staining of Whole Adipose

Mice were euthanized, whole subcutaneous white adipose tissue (scWAT) depots were carefully removed to remain intact and fixed in 2% PFA at 4°C for 4hr-12hrs depending on thickness of tissue. The tissues were then rinsed for 10 minutes with 1X PBS, twice at 4°C. Tissues were stained with Luxol Blue Myelin stain kit (Abcam, Cambridge, U.K. Cat. #ab150675). Tissues were incubated in Luxol Fast Blue Solution overnight at room temperature, briefly rinsed with distilled water and differentiated in Lithium Carbonate Solution followed by 70% Alcohol Reagent until the solution ran clear and adipocytes were colorless while nerves remained blue. Tissue was then rinsed in distilled water, mounted, and imaged on a dissecting microscope.

### Optical Clearing

Optical clearing methods were tested in inguinal scWAT and/or BAT from either *TH-Cre x ROSA YFP* (Jackson Laboratory, Bar Harbor, ME, USA; Stock # 008601 and 006148) reporter mice or *C57BL/6J* mice (The Jackson Laboratory, Stock # 000664). ScaleA2 method described by Hama, H. et al. [34]; sucrose gradient method as described by Tsai, P. et al. [66]; UbasM method as described by Chen, L. et al. [40]. BABB method as described by Li, L. et al. [36]; CUBIC and CUBIC CB-perfusion as described by Susaki, E. et al. [37]; uDISCO as described by Pan, C. et al. [39]; iDISCO as described by Reinier, N. et al. [38]

### Cell Size Quantification

Whole adipose depots were excised from male C57BL/6J mice and fixed in 2% PFA overnight. Whole tissues were either z-depth reduced and mounted on a glass slide (z-depth reduced) or remained unmanipulated and placed on a glass slide (control). Representative images were taken of each tissue and cells with clear in-focus boundaries were outlined. Cell area was measured in FIJI [65].

### Hematoxylin Staining

Whole inguinal adipose depots were excised from male C57BL/6J, fixed in 10% buffered formalin overnight and paraffin embedded. Paraffin embedded tissues were sectioned at 7μm and stained with hematoxylin to increase cell membrane contrast. Tissues were then cover slipped, sealed, and imaged.

### Tissue Sectioning and IF for lysm cre Tissues

Both axillary and inguinal scWAT were collected from adult 7-day cold (5°C) exposed LysM-Cre-Rosa eYFP reporter mice. Whole depots were cut into 100 μm-thick slices longitudinally (from anterior to posterior). Slices from each depot were fixed in 2% PFA for 1hr, then rinsed for 10 minutes with 1X PBS w/ 10U/mL heparin, twice at 4°C. Tissues were incubated in blocking buffer (1XPBS/2.5% BSA/0.5-1% Triton) at 4°C overnight. Tissues were rinsed in rinse buffer and stained with TrueBlack®. Tissue slices were stained with the primary antibody, PSD95 (ab18258 at 1:1000) for 48 hours. Then they were stained with goat anti-rabbit secondary antibody bound to AlexaFluor 647 (1:1000) overnight. Finally, they were stained with the primary antibody conjugated to GFP (ab6662 1:1000) for 72 hours. Sections underwent z-depth reduction as previously describes in Methods Section prior to mounting. Tissues were either imaged on a confocal microscope at 63X magnification. All tissues imaged with a confocal microscope were stained with DAPI in addition to PSD95, AlexaFluor 647 and GFP.

### Transmission Electron Microscopy

Contralateral inguinal and axillary tissue samples were collected as 1 mm^3^ blocks. Tissues were fixed in 2% glutaraldehyde-2% paraformaldehyde for 2 hours and then washed overnight in t0.1 M PB (pH 7.4) buffer at 4°C. Samples were then post-fixed in 1% OsO^4^ in PB, dehydrated in an acetone series (50%, 70%, 95% and 100%) and embedded and polymerized at 60 °C for 48 hrs in Epon/Araldite resin. Once the tissue samples were in resin, an ultramicrotome (Leica UC6) was used to cut the tissues into sections that did not exceed 60-90 nm in thickness. Those sections were then collected directly on a 300-mesh nickel grid(s) and stained with uranyl acetate and lead citrate. The samples were imaged using a CM10 transmission electron microscope (Philips). Images were taken of both adipocytes and cells within the SVF.

### Second Harmonic Generation Imaging

All 2-photon microscopy studies used a modified Olympus FV300 system with an upright BX50WI microscopy stand (Olympus, Center Valley, Pennsylvania) and a mode-locked Ti:Sapphire laser (Chameleon, Coherent, Santa Clara, California). Laser power was modulated via an electro-optic modulator (ConOptics, Danbury, Connecticut). The fluorescence and SHG signals were collected in a non-descanned geometry using a single PMT (H7422 GaAsP, Hamamatsu, Hamamastu City, Japan). Emission wavelengths were separated from excitation wavelengths using a 665 nm dichoric beamsplitter followed by 582/64 nm and 448/20 nm bandpass filters for Alexa 488 and SHG signals respectively (Semrock, Rochester, New York). Images were acquired using circular polarization with excitation power ranging from 1-50 mW and a 40X (0.8 NA) water immersion objective with 3X optical zoom with scanning speeds of 2.71s/frame. All images were 515 × 512 pixels with a field of view of 85 μm.

### Image Processing and Figure Formatting

Images were compiled and formatted for figures in PowerPoint (Microsoft.) In some cases, brightness was increased to enhance small structure visibility. Figure brightness was adjusted identically when direct comparisons were made. Figures were each exported as a Tiff (1650×2200) and then reduced to 300×600 DPI and 8-bit depth for publication.

## Resource Availability

### Lead Contact

Further information and requests for resources and reagents should be directed to and will be fulfilled by the Lead Contact, Kristy Townsend.

### Materials Availability

This study did not generate new unique reagents.

### Data and Code Availability

Neurite density quantification and heat map generating code developed for this publication can be found on GitHub at https://github.com/ktownsendlab/willows_et_al-2020. Protocols used for this publication can be found on Protocols.io at https://dx.doi.org/10.17504/protocols.io.brs2m6ge. All other data is included in this manuscript.

## Supplemental Information

Supplemental Table S1: Comparison of methods for clearing adipose tissue

Supplemental Figure S1: Resonant scanning as an alternative imaging approach.

Supplemental Figure S2: Additional whole mount staining optimization.

Supplemental Figure S3: Height color coded z-maximum projection.

Supplemental Figure S4: 7μm cross section of peripheral nerve bundles in inguinal scWAT.

Supplemental Figure S5: Transmission Electron Microscopy (TEM) of scWAT Innervation.

Supplemental Figure S6: Synapsing within white adipose tissue.

Supplemental Figure S7: Additional nerve labeling in scWAT.

Supplemental Figure S8: Measuring neurite density from z-maximum projections is equivalent to measuring from individual z-slices.

## References Cited

1. Blaszkiewicz, M., et al., The Importance of Peripheral Nerves in Adipose Tissue for the Regulation of Energy Balance. Biology (Basel), 2019. 8(1).

2. Wirsen, C., Adrenergic Innervation of Adipose Tissue Examined by Fluorescence Microscopy. Nature, 1964. 202: p. 913.

3. Cinti, S., et al., Tim Bartness, Ph.D. (1953-2015). Temperature (Austin), 2016. 3(1): p. 31–8.

4. Fenzl, A. and F.W. Kiefer, Brown adipose tissue and thermogenesis. Horm Mol Biol Clin Investig, 2014. 19(1): p. 25–37.

5. Bartness, T.J., C.H. Vaughan, and C.K. Song, Sympathetic and sensory innervation of brown adipose tissue. Int J Obes (Lond). 2010. 34 Suppl 1:S36-42.: p. S36–S42.

6. Francois, M., et al., Sympathetic innervation of the interscapular brown adipose tissue in mouse. Ann N Y Acad Sci, 2019.

7. Garretson, J.T., et al., Lipolysis sensation by white fat afferent nerves triggers brown fat thermogenesis. Molecular metabolism, 2016. 5(8): p. 626–634.

8. Foster, D.O., F. Depocas, and M. Zuker, Heterogeneity of the sympathetic innervation of rat interscapular brown adipose tissue via intercostal nerves. Can J Physiol Pharmacol, 1982. 60(6): p. 747–54.

9. Foster, M.T. and T.J. Bartness, Sympathetic but not sensory denervation stimulates white adipocyte proliferation. Am J Physiol Regul Integr Comp Physiol, 2006. 291(6): p. R1630–7.

10. Fishman, R.B. and J. Dark, Sensory innervation of white adipose tissue. Am J Physiol, 1987. 253(6 Pt 2): p. R942–4.

11. Giordano, A., et al., White adipose tissue lacks significant vagal innervation and immunohistochemical evidence of parasympathetic innervation. Am J Physiol Regul Integr Comp Physiol, 2006. 291(5): p. R1243–55.

12. Murano, I., et al., Noradrenergic parenchymal nerve fiber branching after cold acclimatisation correlates with brown adipocyte density in mouse adipose organ. J Anat, 2009. 214(1): p. 171–8.

13. Vargovic, P., et al., Adipocytes as a new source of catecholamine production. FEBS.Lett., 2011. 585(14): p. 2279–2284.

14. Shi, H., et al., Sensory or sympathetic white adipose tissue denervation differentially affects depot growth and cellularity. Am J Physiol Regul Integr Comp Physiol, 2005. 288(4): p. R1028–37.

15. Blaszkiewicz, M., et al., Neuropathy and neural plasticity in the subcutaneous white adipose depot. PLoS One, 2019. 14(9): p. e0221766.

16. Dichamp, J., et al., 3D analysis of the whole subcutaneous adipose tissue reveals a complex spatial network of interconnected lobules with heterogeneous browning ability. Sci Rep, 2019. 9(1): p. 6684.

17. Zeng, W., et al., Sympathetic neuro-adipose connections mediate leptin-driven lipolysis. Cell, 2015. 163(1): p. 84–94.

18. Cao, Y., et al., Three-dimensional volume fluorescence-imaging of vascular plasticity in adipose tissues. Mol Metab, 2018. 14: p. 71–81.

19. Cao, Y., H. Wang, and W. Zeng, Whole-tissue 3D imaging reveals intra-adipose sympathetic plasticity regulated by NGF-TrkA signal in cold-induced beiging. Protein Cell, 2018. 9(6): p. 527–539.

20. Chi, J., et al., Three-Dimensional Adipose Tissue Imaging Reveals Regional Variation in Beige Fat Biogenesis and PRDM16-Dependent Sympathetic Neurite Density. Cell Metab, 2018. 27(1): p. 226–236 e3.

21. Li, X., et al., Co-staining Blood Vessels and Nerve Fibers in Adipose Tissue. J Vis Exp, 2019(144).

22. Jiang, H., et al., Dense Intra-adipose Sympathetic Arborizations Are Essential for Cold-Induced Beiging of Mouse White Adipose Tissue. Cell Metab, 2017. 26(4): p. 686–692 e3.

23. Kim, S.Y., et al., Stochastic electrotransport selectively enhances the transport of highly electromobile molecules. Proc Natl Acad Sci U S A, 2015. 112(46): p. E6274–83.

24. Fiorelli, R., et al., Enhanced tissue penetration of antibodies through pressurized immunohistochemistry. bioRxiv, 2020: p. 2020.09.25.311936.

25. Kim, J.Y., et al., BrainFilm, a novel technique for physical compression of 3D brain slices for efficient image acquisition and post-processing. Sci Rep, 2018. 8(1): p. 8531.

26. Thompson, R.J., et al., PGP 9.5--a new marker for vertebrate neurons and neuroendocrine cells. Brain Res, 1983. 278(1-2): p. 224–8.

27. Day, I.N. and R.J. Thompson, UCHL1 (PGP 9.5): neuronal biomarker and ubiquitin system protein. Prog Neurobiol, 2010. 90(3): p. 327–62.

28. Draberova, E., et al., Class III beta-tubulin is constitutively coexpressed with glial fibrillary acidic protein and nestin in midgestational human fetal astrocytes: implications for phenotypic identity. J Neuropathol Exp Neurol, 2008. 67(4): p. 341–54.

29. Latremoliere, A., et al., Neuronal-Specific TUBB3 Is Not Required for Normal Neuronal Function but Is Essential for Timely Axon Regeneration. Cell Rep, 2018. 24(7): p. 1865–1879 e9.

30. Hasegawa, H., et al., Analyzing somatosensory axon projections with the sensory neuron-specific Advillin gene. J Neurosci, 2007. 27(52): p. 14404–14.

31. Hunter, D.V., et al., Advillin Is Expressed in All Adult Neural Crest-Derived Neurons. eNeuro, 2018. 5(5).

32. Bird, E.V., et al., Correlation of Nav1.8 and Nav1.9 sodium channel expression with neuropathic pain in human subjects with lingual nerve neuromas. Mol Pain, 2013. 9: p. 52.

33. D’Urso, D., et al., Protein zero of peripheral nerve myelin: biosynthesis, membrane insertion, and evidence for homotypic interaction. Neuron, 1990. 4(3): p. 449–60.

34. Hama, H., et al., Scale: a chemical approach for fluorescence imaging and reconstruction of transparent mouse brain. Nat Neurosci, 2011. 14(11): p. 1481–8.

35. Dodt, H.U., et al., Ultramicroscopy: three-dimensional visualization of neuronal networks in the whole mouse brain. Nat Methods, 2007. 4(4): p. 331–6.

36. Li, L., et al., The functional organization of cutaneous low-threshold mechanosensory neurons. Cell, 2011. 147(7): p. 1615–27.

37. Susaki, E.A., et al., Advanced CUBIC protocols for whole-brain and whole-body clearing and imaging. Nat Protoc, 2015. 10(11): p. 1709–27.

38. Renier, N., et al., iDISCO: a simple, rapid method to immunolabel large tissue samples for volume imaging. Cell, 2014. 159(4): p. 896–910.

39. Pan, C., et al., Shrinkage-mediated imaging of entire organs and organisms using uDISCO. Nat Methods, 2016. 13(10): p. 859–67.

40. Chen, L., et al., UbasM: An effective balanced optical clearing method for intact biomedical imaging. Sci Rep, 2017. 7(1): p. 12218.

41. Brantschen, S., et al., Regulatory effect of recombinant interleukin (IL)3 and IL4 on cytokine gene expression of bone marrow and peripheral blood mononuclear cells. European Journal of Immunology, 1989. 19: p. 2017–2023.

42. Azaripour, A., et al., A survey of clearing techniques for 3D imaging of tissues with special reference to connective tissue. Prog Histochem Cytochem, 2016. 51(2): p. 9–23.

43. Matryba, P., L. Kaczmarek, and J. Gołąb, Advances in Ex Situ Tissue Optical Clearing. Laser & Photonics Reviews, 2019. 13(8): p. 1800292.

44. Yu, T., et al., Optical clearing for multiscale biological tissues. J Biophotonics, 2018. 11(2).

45. Seo, J., M. Choe, and S.Y. Kim, Clearing and Labeling Techniques for Large-Scale Biological Tissues. Mol Cells, 2016. 39(6): p. 439–46.

46. Croce, A.C. and G. Bottiroli, Autofluorescence spectroscopy and imaging: a tool for biomedical research and diagnosis. Eur J Histochem, 2014. 58(4): p. 2461.

47. Schnell, S.A., W.A. Staines, and M.W. Wessendorf, Reduction of Lipofuscin-like Autofluorescence in Fluorescently Labeled Tissue. Journal of Histochemistry & Cytochemistry, 1999. 47(6): p. 719–730.

48. Song, E., et al., Optical clearing based cellular-level 3D visualization of intact lymph node cortex. Biomed Opt Express, 2015. 6(10): p. 4154–64.

49. Chung, K., et al., Structural and molecular interrogation of intact biological systems. Nature, 2013. 497(7449): p. 332–7.

50. Stradleigh, T.W., et al., Moniliform deformation of retinal ganglion cells by formaldehyde-based fixatives. J Comp Neurol, 2015. 523(4): p. 545–64.

51. Bartness, T.J. and M. Bamshad, Innervation of mammalian white adipose tissue: implications for the regulation of total body fat. Am J Physiol, 1998. 275(5): p. R1399–411.

52. Sheng, Y. and L. Zhu, The crosstalk between autonomic nervous system and blood vessels. Int J Physiol Pathophysiol Pharmacol, 2018. 10(1): p. 17–28.

53. Martinez-Santibanez, G., K.W. Cho, and C.N. Lumeng, Imaging white adipose tissue with confocal microscopy. Methods Enzymol, 2014. 537: p. 17–30.

54. Ernst, C. and B.R. Christie, Isolectin-IB 4 as a vascular stain for the study of adult neurogenesis. J Neurosci Methods, 2006. 150(1): p. 138–42.

55. Peters, B.P. and I.J. Goldstein, The use of fluorescein-conjugated Bandeiraea simplicifolia B4-isolectin as a histochemical reagent for the detection of alpha-D-galactopyranosyl groups. Their occurrence in basement membranes. Exp Cell Res, 1979. 120(2): p. 321–34.

56. Ismail, J.A., et al., Immunohistologic labeling of murine endothelium. Cardiovasc Pathol, 2003. 12(2): p. 82–90.

57. Gorakshakar, A.C. and K. Ghosh, Use of lectins in immunohematology. Asian J Transfus Sci, 2016. 10(1): p. 12–21.

58. Rechthand, E., et al., Distribution of adrenergic innervation of blood vessels in peripheral nerve. Brain Res, 1986. 374(1): p. 185–9.

59. Klabunde, R.E., Cardiovascular physiology concepts. 2nd ed. 2012, Philadelphia, PA: Lippincott Williams & Wilkins/Wolters Kluwer. xi, 243 p.

60. Thomas, G.D., Neural control of the circulation. Adv Physiol Educ, 2011. 35(1): p. 28–32.

61. Westcott, E.B. and S.S. Segal, Perivascular innervation: a multiplicity of roles in vasomotor control and myoendothelial signaling. Microcirculation, 2013. 20(3): p. 217–38.

62. Müller, B., et al., High-resolution tomographic imaging of microvessels. Proc SPIE, 2008. 7078: p. 70780B.

63. Fang, X., et al., Intense isolectin-B4 binding in rat dorsal root ganglion neurons distinguishes C-fiber nociceptors with broad action potentials and high Nav1.9 expression. J Neurosci, 2006. 26(27): p. 7281–92.

64. Vulchanova, L., et al., Cytotoxic targeting of isolectin IB4-binding sensory neurons. Neuroscience, 2001. 108(1): p. 143–55.

65. Schindelin, J., et al., Fiji: an open-source platform for biological-image analysis. Nat Methods, 2012. 9(7): p. 676–82.

66. Tsai, P.S., et al., Correlations of neuronal and microvascular densities in murine cortex revealed by direct counting and colocalization of nuclei and vessels. J Neurosci, 2009. 29(46): p. 14553–70.

