## Supplemental Figures for "Visualization and Analysis of Whole Depot Adipose Tissue Neural Innervation"

### Supplemental Table S1: Comparison of methods for clearing adipose tissue

| | Level of Clearing for Adipose | Labeling (immunofluorescence (IF), endogenous fluorescent protein) | Tissue Morphological Change | Corrosive to Lenses? |
|---|---|---|---|---|
| **ScaleA2** 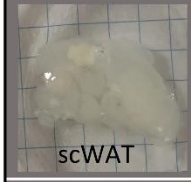  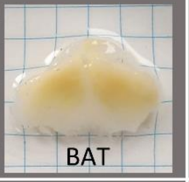  | Poor | Not ideal for IF<br>Good for endogenous signal | Tissue expansion | No |
| **Sucrose** 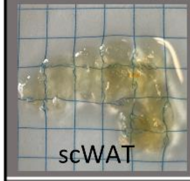  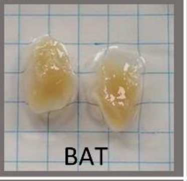  | Mediocre | Good for both | Tissue Shrinkage | No |
| **CUBIC** 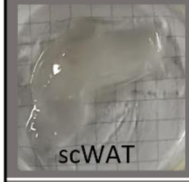  | Mediocre | Good for IF<br>Some quenching of endogenous signal | Tissue expansion | No |
| **CUBIC CB-Perfusion** 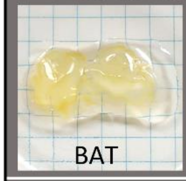  | Mediocre | Good for IF<br>Good with endogenous signal | Tissue expansion; foam-like | No |
| **UbasM** 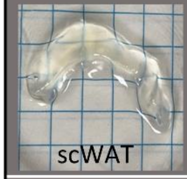  | Good | Good for both | Tissue expansion | No |
| **BABB** 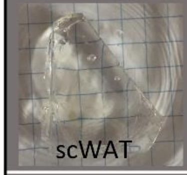  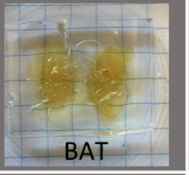  | Excellent | Excellent for IF<br>Endogenous signals require immunolabeling | Tissue shrinkage; increased rigidity | Yes (BABB) |
| **iDISCO** 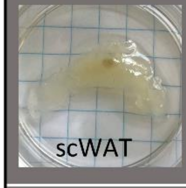  | Excellent | Excellent for IF<br>Endogenous signals require immunolabeling | Tissue shrinkage, discoloration, and increased rigidity | Yes (DME) |
| **uDISCO** 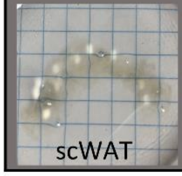  | Excellent | Excellent for IF<br>Endogenous signals require immunolabeling | Tissue shrinkage, discoloration, and increased rigidity | Yes (DME) |

Supplemental Figure S1: Resonant scanning as an alternative imaging approach

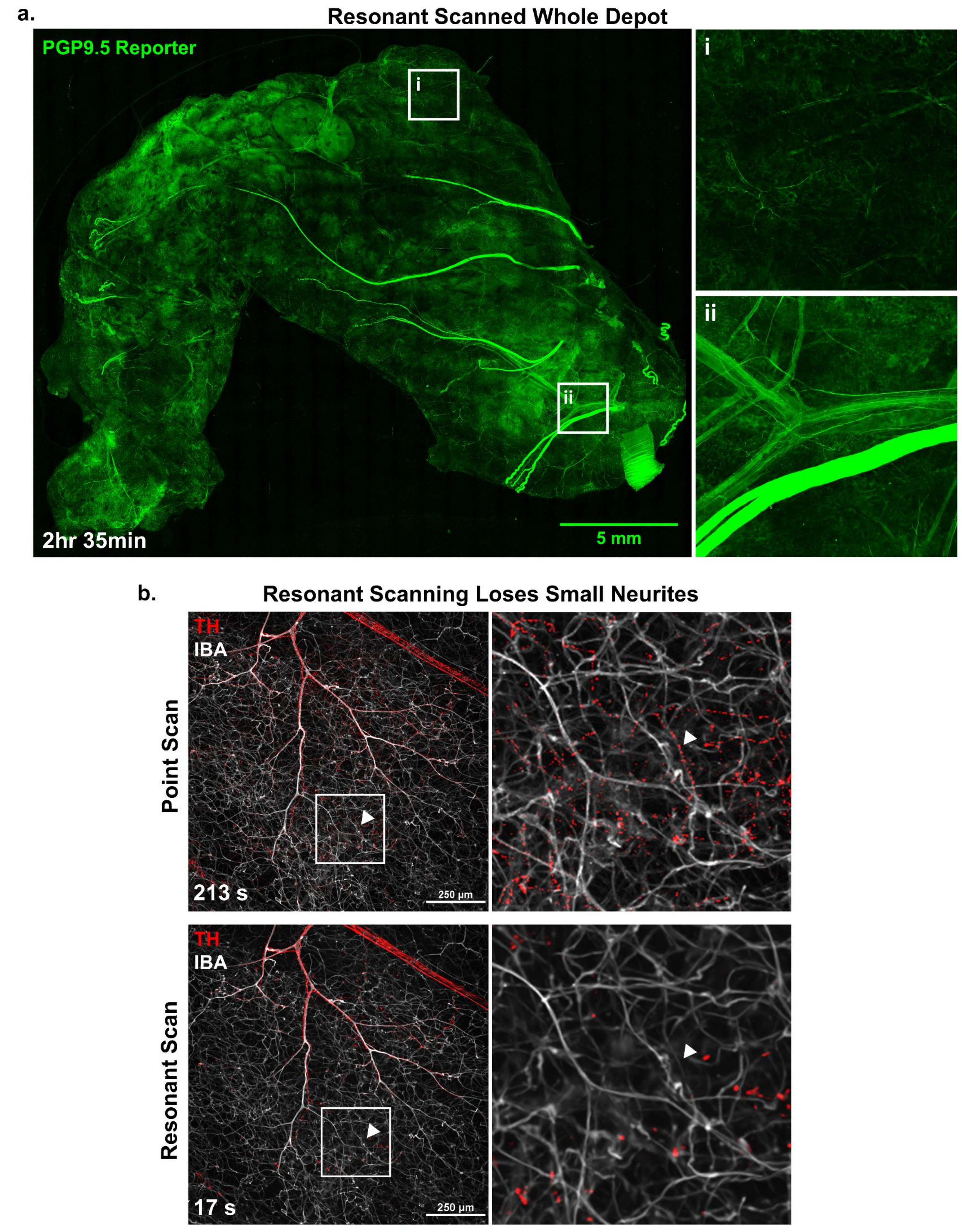

### Supplemental Figure S2: Additional whole mount staining optimization

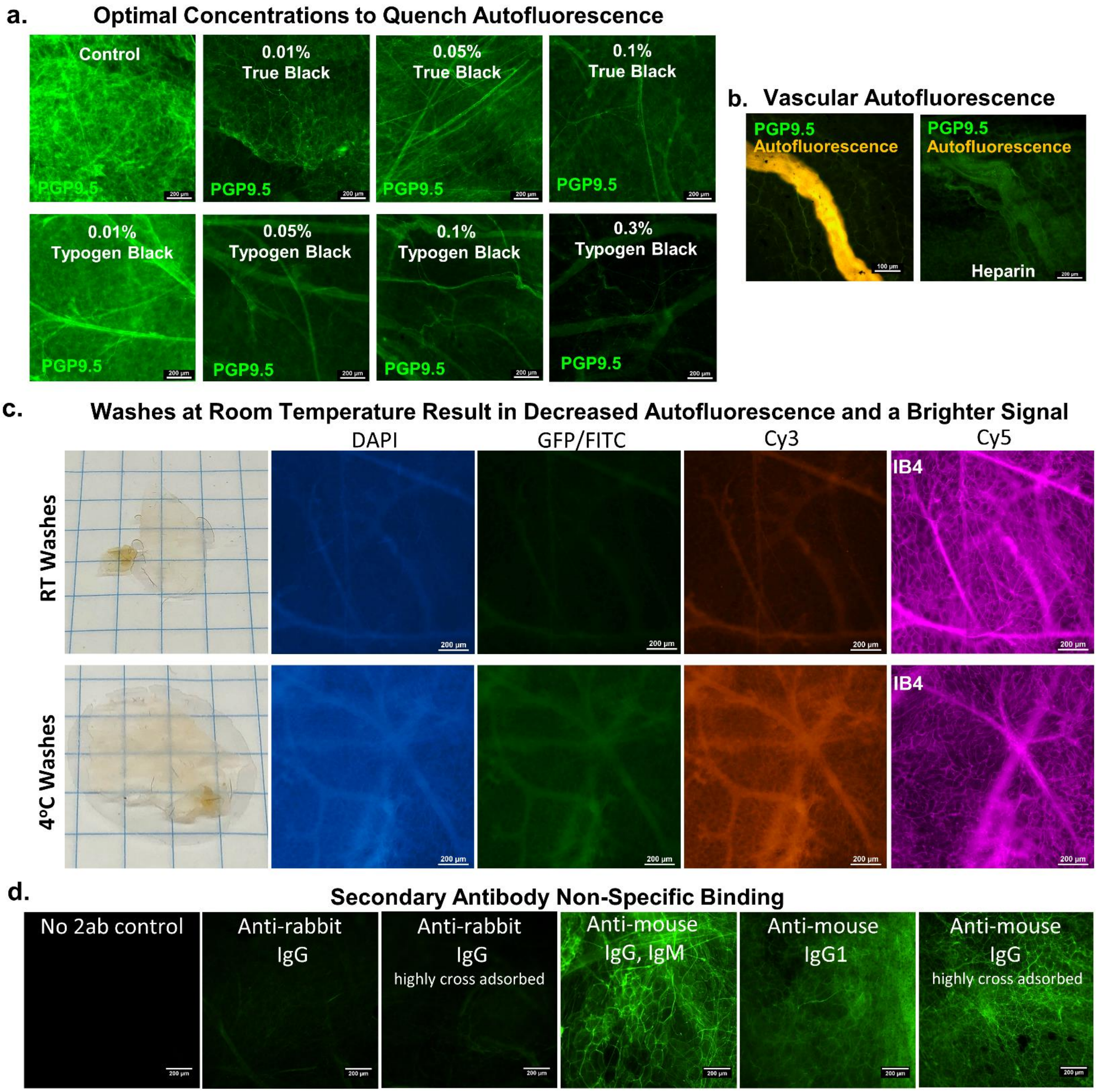

### Supplemental Figure S3: Height color coded z-maximum projection

**a.**

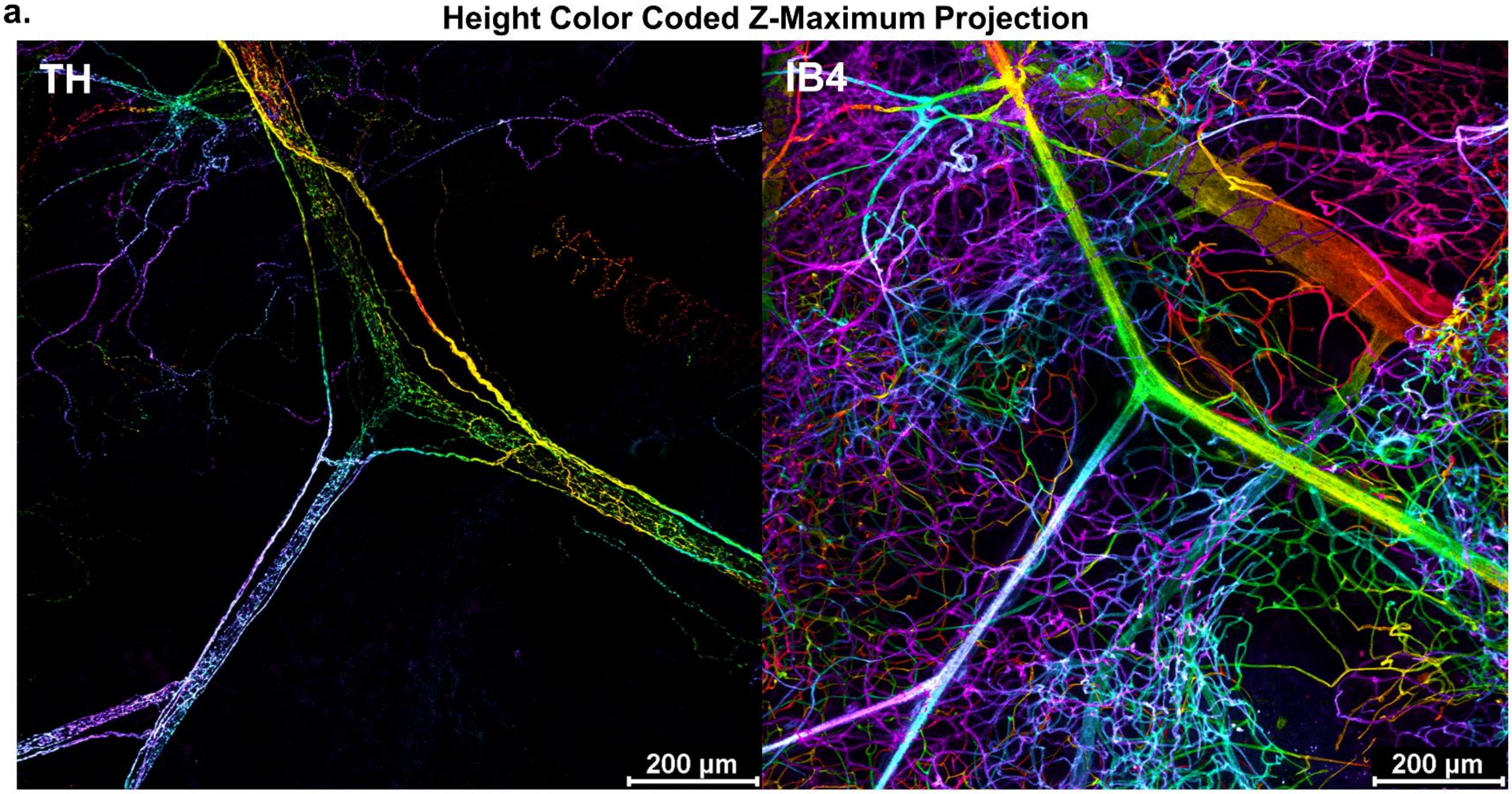

**Supplemental Figure S4:** 7μm cross section of peripheral nerve bundles in inguinal scWAT

**a.** 7μm cross section of peripheral nerve bundles in inguinal scWAT

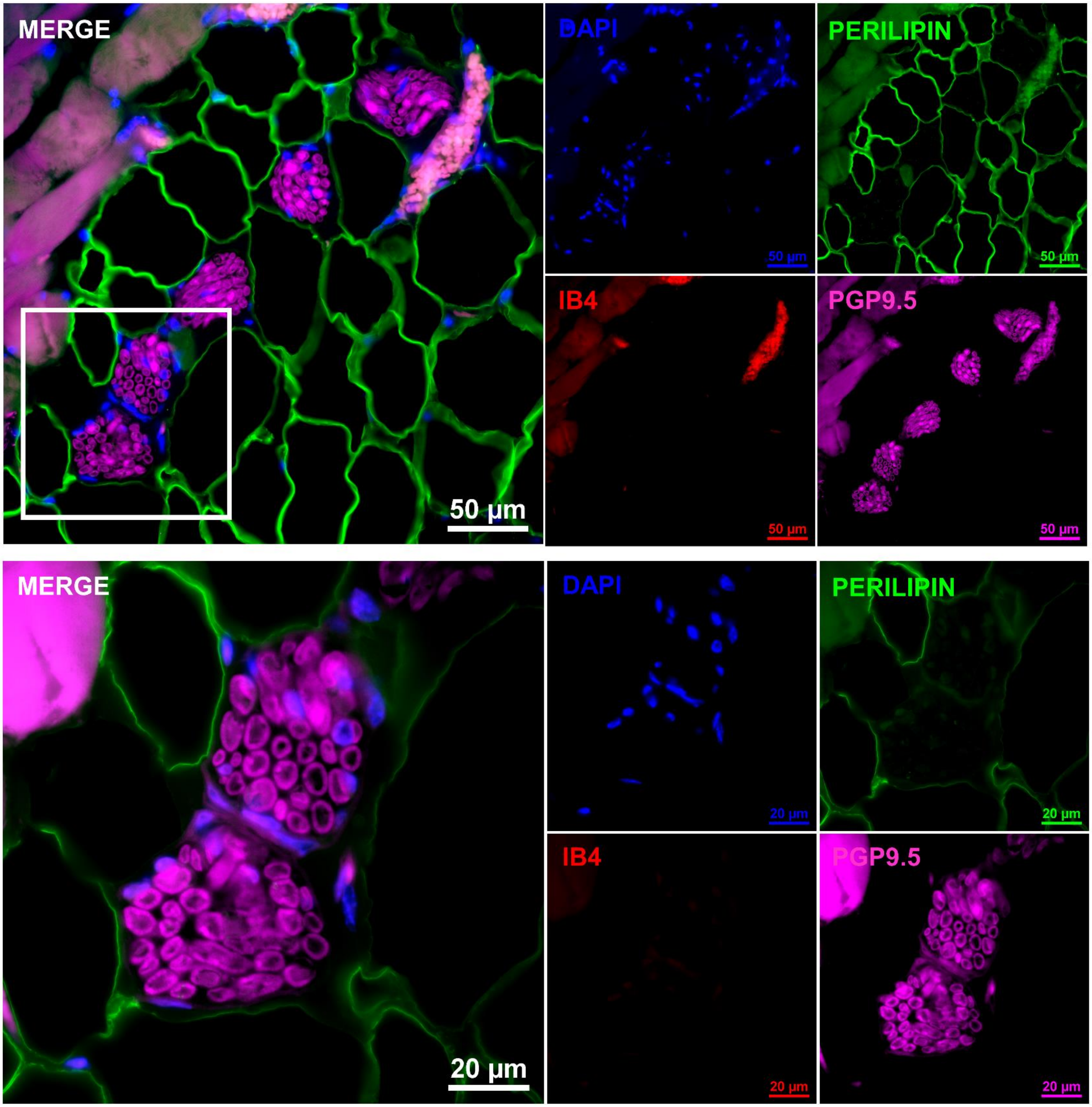

### Supplemental Figure S5: Transmission electron microscopy of scWAT innervation

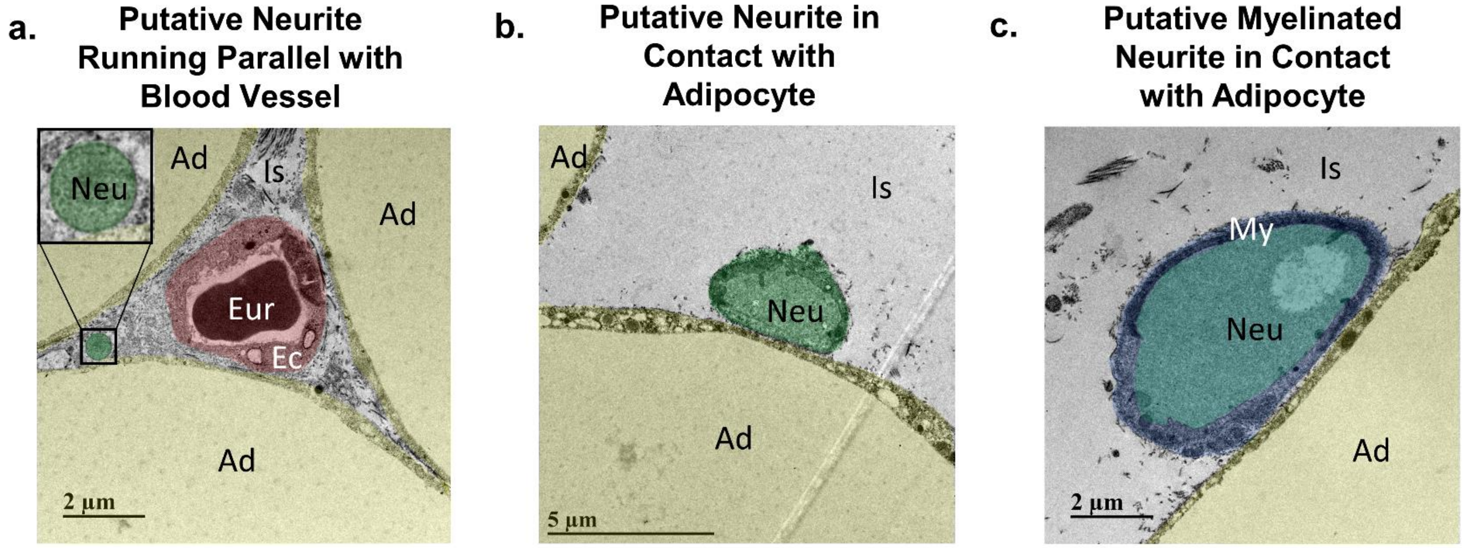

### Supplemental Figure S6: Synapsing within white adipose tissue

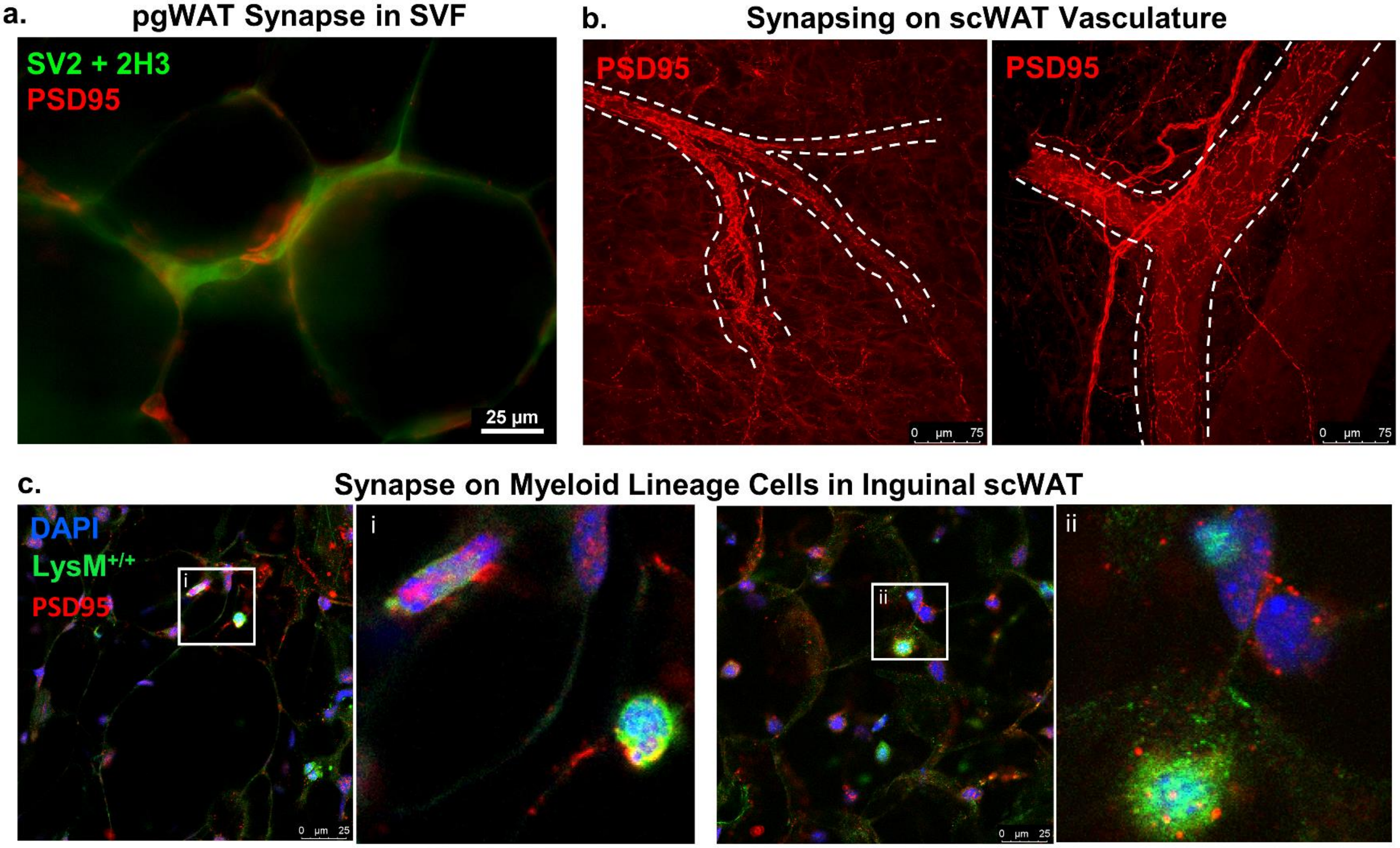

### Supplemental Figure S7: Additional nerve labeling in scWAT

**a.** **Sensory Nerve Reporter in Inguinal scWAT**

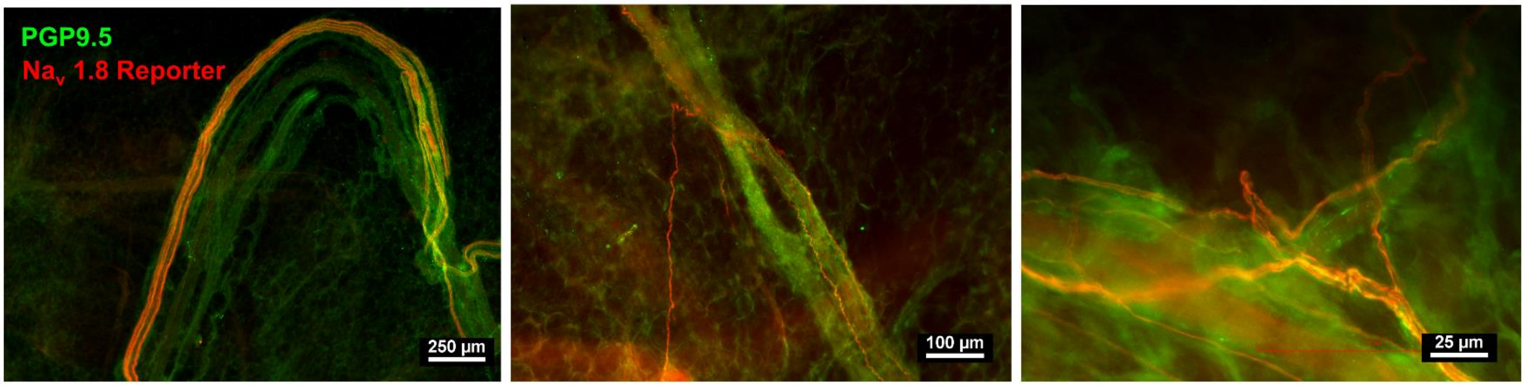


**b.** **Inguinal scWAT Luxol Blue Myelin Stain**

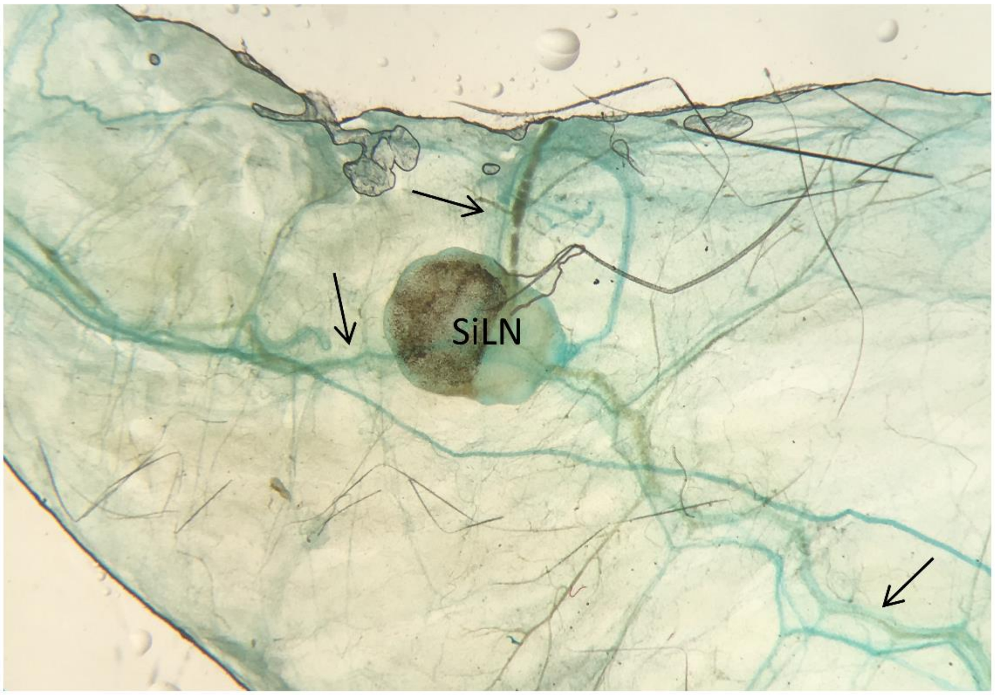

**Supplemental Figure S8: Measuring neurite density from z-maximum projections is equivalent to measuring from individual z-slices**

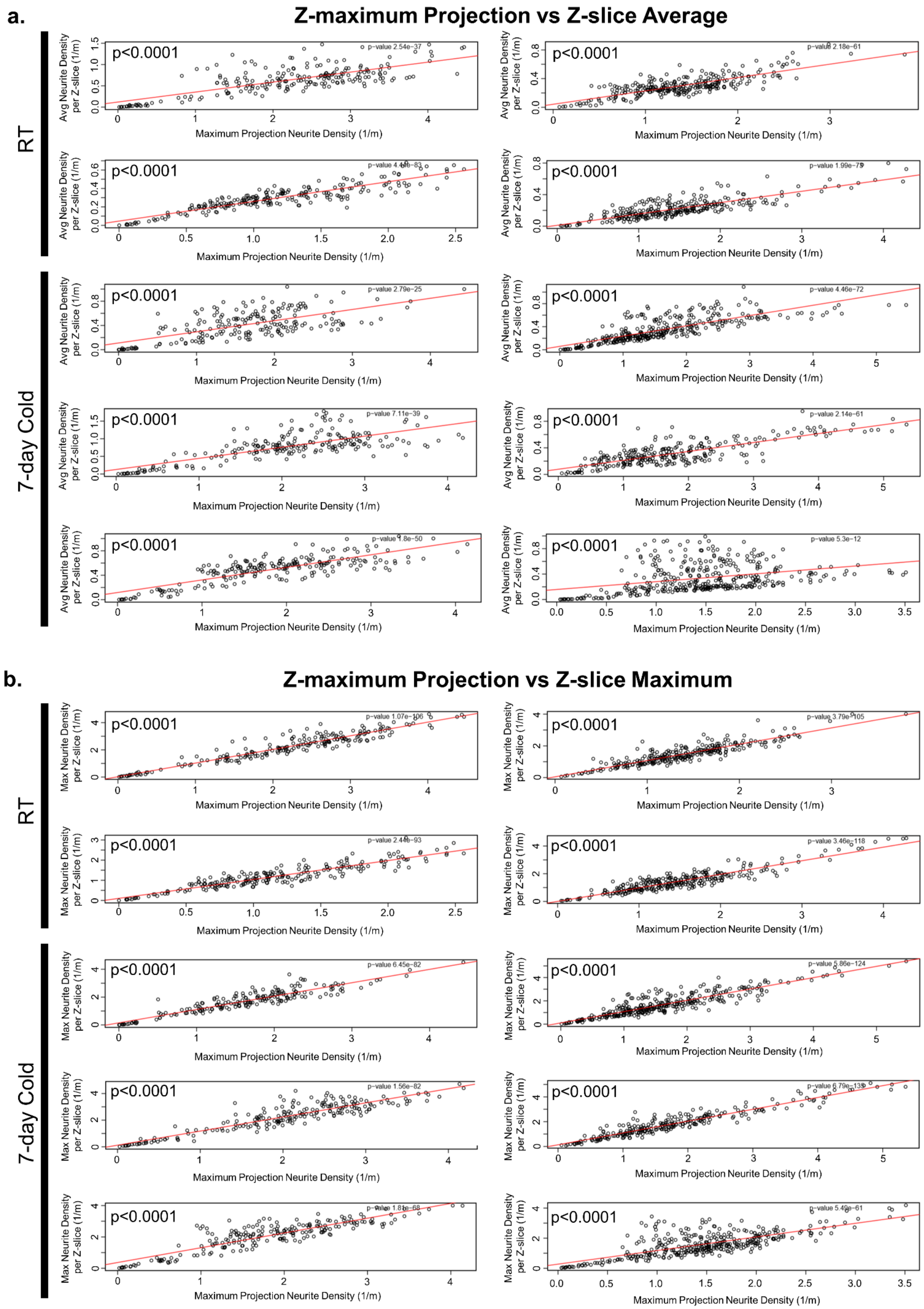

**Supplemental Figure S9: Second harmonic generation imaging of collagen in BTBR$^{+/+}$ (WT) and BTBR$^{ob/ob}$ (MUT) inguinal scWAT**

**a.** **Second Harmonic Generation Imaging of Collagen in BTBR Inguinal scWAT**

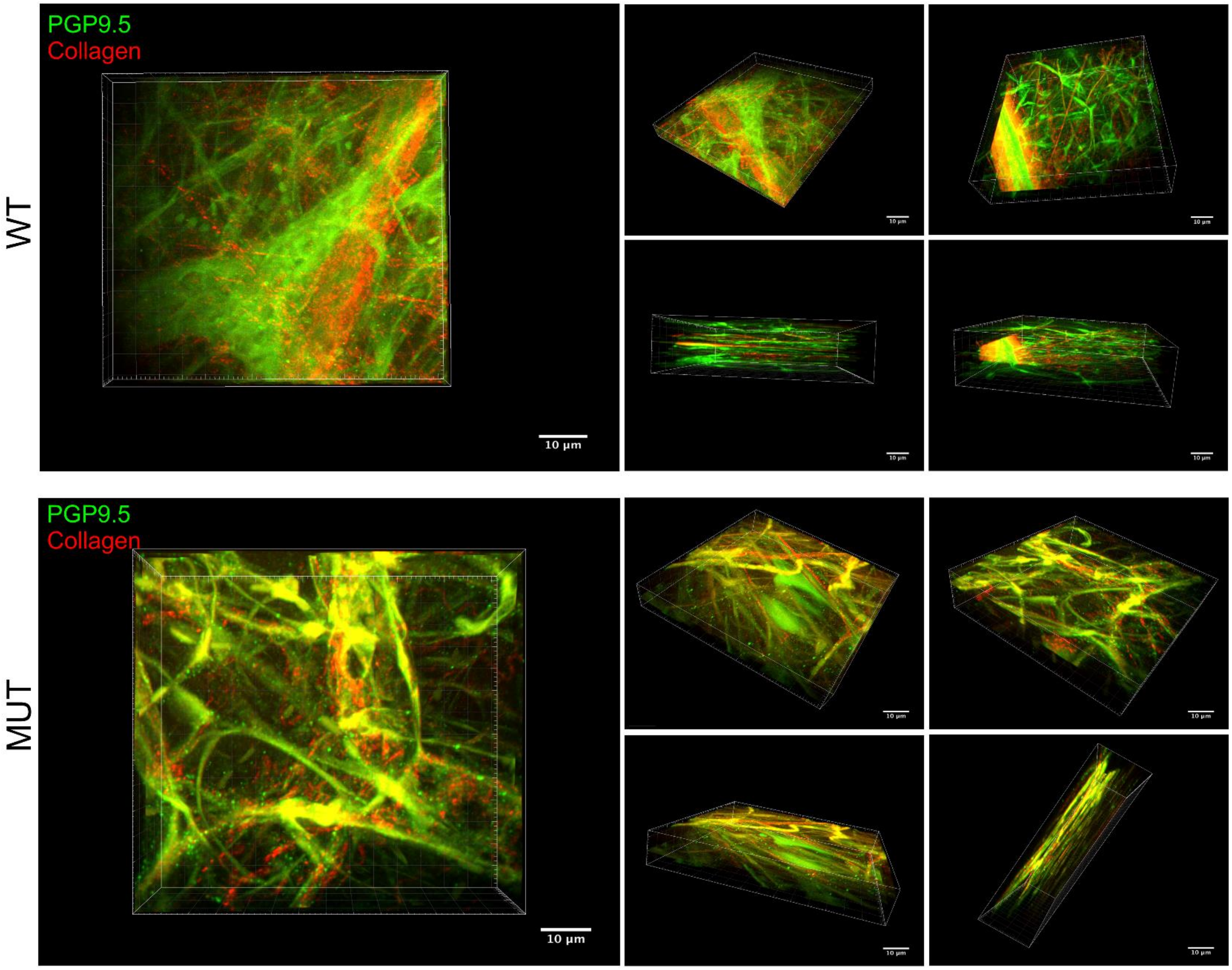
