## Supplemental Text for "Visualization and Analysis of Whole Depot Adipose Tissue Neural Innervation"

Supplemental Figure Legends

**Supplemental Table S1: Comparison of methods for clearing adipose tissue**

**Supplemental Figure S1: Resonant scanning as an alternative imaging approach.** Intact inguinal subcutaneous white adipose tissue (scWAT) depot was excised from a male pan-neuronal direct reporter mouse (*C57BL/6-Tg(Uchl1-EGFP)G1Phoz/J*) and whole mount processed. Tissue was resonant scanned with a 10X objective at 1024 x 1024 resolution, with a Z-step size of 16 μm and was tiled and Z-maximum projected (28x23x19; 12,236 images) and denoised using (a). Total time to image was only 2 hours and 35 minutes (a). White boxes were digitally zoomed by 5.35X (i-ii). Inguinal depot from male *C57BL/6J* mouse was whole mount processed and immunostained with Tyrosine Hydroxylase (TH) in red and Isolectin $IB_4$ (IB4) in white. The same area and thickness (90.43 μm) of tissue was imaged with a 6 μm step size at 1024 x 1024 resolution and scanned unidirectionally with both point scanning (213 seconds) and resonant scanning (17 seconds) (b). 4.43X digital zoom-ins help illustrate the presence or absence of small TH+ neurites. The same example is marked by the white arrow (b). All images were captured using Nikon A1R Confocal microscope and resonant scanned images were denoised with Nikon Elements Denoise.ai (a-b). Scale bars are 5 mm (a) 250 μm (b).

**Supplemental Figure S2: Additional whole mount staining optimization.** Autofluorescence quenching dilutions were compared to an unquenched control tissue in conjunction with pan-neuronal PGP9.5 staining (green) to identify what concentrations allowed for highest nerve visibility (a). Whole ingunal scWAT depots received either washes with 1XPBS or 1XPBS w/ 10 U/mL Heparin. Tissues were then immunostained with the pan-neuronal marker PGP9.5 (green.) Blood vessel autofluorescence (orange) was significantly reduced when tissues were washed with 1XPBS/10U/ml Heparin (b). Imaged at 20X with (DS-fi2) and at 10X imaged with (ORCA-

Flash4.0 V2) (b). Axillary scWAT was whole mount processed and one tissue received all washes at room temperature and the other received washes at 4ºC. Tissues were stained with the vasculature marker Isolectin $IB_4$ (IB4) conjugated to 647nm fluorophore and imaged with 4 filter cubes: DAPI, GFP/FITC, Cy3, and Cy5 (c). Left-most images demonstrate relative tissue clarity of whole depot. DAPI, GFP/FITC, and Cy3 images demonstrate relative autofluorescence in each channel. Cy5 image demonstrates relative IB4 fluorescence signal intensity. 10X objective (c). Comparison of non-specific secondary antibody binding in whole mount processed inguinal scWAT. Secondary antibodies were conjugated to 488nm fluorophores. Captured with 10X objective (d). All micrographs were captured on Nikon E400 microscope (a-d). Scale bars are 200 µm (a,c-d) 100 µm and 200 µm (b).

**Supplemental Figure S3: Height color coded z-maximum projection.** Inguinal scWAT depot was excised from mouse and whole mount processed. Tissue was immunostained with TH and IB4. Imaged as a z-maximum projection and height color coded in LASX. Micrograph captured on Leica TCS SP8 DLS confocal microscope with 10X objective (a). Purple (foreground), green (middle), red (background) (a). Scale bars are 200 µm.

**Supplemental Figure S4: 7µm cross section of peripheral nerve bundles in inguinal scWAT.** Axillary scWAT excised from mouse was embedded in paraffin, sectioned at 7 µm thick, and stained with DAPI (blue), perilipin (green), IB4 (red), PGP9.5 (purple). Micrograph of 5 nerve bundles running parallel captured with 40X objective (upper panels) and 100X objective (lower panels). Captured on Nikon E400 epifluorescence microscope. Scale bars are 50 µm and 20 µm.

**Supplemental Figure S5: Transmission Electron Microscopy (TEM) of scWAT Innervation.** TEM micrographs of putative neurites residing within inguinal scWAT (a-c). Putative neurites in contact with adipocytes and near a blood vessel (a) unmyelinated (b) and myelinated (c).

Captured on CM10 transmission electron microscope at 5800X (a,c) and 4600X (b). Adipocyte (Ad), interstitial space (Is), myelin (My), neuron (Neu), eukaryote (Eur); endothelial cell (Ec). Scale bars are 2 μm (a,c) and 5 μm (b).

**Supplemental Figure S6: Synapsing within white adipose tissue.** Perigonadal white adipose tissue (pgWAT) depot from *C57BL/6J* mouse was whole mount processed and immunostained with markers for neurofilament (2H3, green) and synaptic vesicles (SV2, green) to illustrate the pre-synaptic axon terminal and the post-synaptic marker PSD95 (red). Imaged with 40X objective lens on Nikon E400 microscope (a). Inguinal scWAT depot immunostained with PSD95 revealed synapsing all along a blood vessel (dashed white border). Imaged with 40X objective lens using extended depth of field (NIS-Elements) on Nikon E400 microscope (b). Synapse on Myeloid lineage cells. Imaged with 63X objective on Leica TCS SP8 DLS microscope, captured as z-max projection (c). White boxes expanded for visualization (4.75X) of PSD9.5+ myeloid lineage cells (c-i, c-ii). Scale bars are 25 μm (a,c) and 75 μm (b).

**Supplemental Figure S7: Additional nerve labeling in scWAT.** Inguinal scWAT depot from a $Na_v$ 1.8 reporter mouse imaged with 4X,10X, and 40X objectives on Nikon E400 microscope (a). Luxol Fast Blue myelin staining of whole inguinal scWAT depot. Image taken on Nikon SMZ800 dissecting microscope at 1X and had a 30X digital zoom applied, arrows point to branches of the thoracoepigastric vein and the subilliac lymph node is labeled as SiLN (b). Inguinal scWAT depot immunostained with Isolectin $IB_4$ (white), β3-Tubulin (green), and advillin (AVIL, red) imaged with 10X objective on TCS SP8 DLS microscope and z-max projected (c). Scale bars, from left to right, are 250 μm, 100 μm, and 25 μm (a).

**Supplemental Figure S8: Measuring neurite density from z-maximum projections is equivalent to measuring from individual z-slices.** Using the data presented in Figure 8a-b; images were z-max projected, tiled, and thresholded and neurite density quantification was performed (as outlined in Figure 1). Alternatively the same analysis was performed without maximum projecting each z-stack and neurite density was calculated for each tile either as an average of all of the neurite density scores across the z-slices (a) or as a maximum of all of the neurite density scores across the z-slices (b) and plotted against the maximum projection neurite densities per respective tile.

**Supplemental Figure S9: Second harmonic generation imaging of collagen in BTBR$^{+/+}$ (WT) and BTBR$^{ob/ob}$ (MUT) inguinal scWAT.** Two-photon microscopy was performed on immunofluorescent stained (PGP9.5) inguinal scWAT of BTBR$^{+/+}$ (WT) and BTBR$^{ob/ob}$ (MUT) animals (a). PGP9.5 was detected by excitation of AlexaFluor 488 at 800nm and emission collected using a 582 +/- 64nm filter. For detection of collagen, samples were excited at 890nm and the SHG signal was collected using a 448 +/-20nm filter. A 40X water immersion objective was used. IMARIS software was used to render 3D projections from z-stacks. Images are representative of N=3 WT/MUT, 12-week-old males. Scale bars are 10 μm.
